# Lactate regulation of activation in CD8+ T cells

**DOI:** 10.1101/2021.12.14.472728

**Authors:** Laura Barbieri, Pedro Veliça, Paulo A. Gameiro, Pedro P. Cunha, Iosifina P. Foskolou, David Bargiela, Helene Rundqvist, Randall S. Johnson

**Affiliations:** Department of Cell and Molecular Biology, Karolinska Institutet, Stockholm, Sweden; Department of Surgery, Oncology, and Gastroenterology, University of Padova, Italy; The Francis Crick Institute, London, UK; Department of Neuromuscular Diseases, UCL Queen, Square Institute of Neurology, London, UK; Department of Physiology, Development and Neuroscience, University of Cambridge, Cambridge, UK; Department of Laboratory Medicine, Karolinska Institutet, Stockholm, Sweden

## Abstract

CD8+ T cells infiltrate virtually every tissue to find and destroy infected or mutated cells. They often traverse varying oxygen levels and nutrient-deprived microenvironments. High glycolytic activity in tissues can result in extended exposure of cytotoxic T cells to the metabolite lactate. Lactate can be immunosuppressive, at least in part due to its association with tissue acidosis. We show here that the lactate anion is well tolerated by CD8+ T cells in pH neutral conditions. We describe how lactate is taken up by activated CD8+ T cells and is capable of displacing glucose as a carbon source. Activation in the presence of a pH neutral form of lactate significantly alters the CD8+ T cell transcriptome, including the expression of key effector differentiation markers such as granzyme B and interferon-gamma. Our studies reveal the novel metabolic features of lactate utilization by activated CD8+ T cells, and highlight the importance of lactate in shaping the differentiation and activity of cytotoxic T cells.

## Introduction

Within the adaptive immune system, cytotoxic CD8+ T cells play a central role in controlling viral infections and tumor growth. To accomplish this, T cells are required to rapidly infiltrate tissues and must therefore be able to adapt to fluctuations in local nutrient levels (Klein Geltink et al., 2018). Upon encountering a cognate antigen, naïve CD8+ T cells undergo a rapid metabolic and transcriptional shift that drives them out of quiescence and into an activated, highly proliferative state (Ashley Menk et al., 2018). This is supported by accumulation of the hypoxia-inducible factor 1 alpha (HIF-1α) leading to increased glucose uptake and breakdown via the glycolytic pathway (Doedens et al., 2013; Palazon et al., 2014; Wang et al., 2011). Rapid ATP production and increased availability of metabolic intermediates, by-products of the glycolytic flux, allow for rapid expansion and reactivity to immunogenic stimuli (Pearce et al., 2013).

Lactate is a major byproduct of glycolysis and its production is considered a marker of T cell activation (Grist et al., 2018). High levels of HIF-1α in turn give rise to striking increases in intracellular lactate following triggering of glycolysis in CD8+ T cells (Tyrakis et al., 2016). Lactate export is coupled with proton release which can result in localized acidosis. Healthy tissue microenvironments, however, are efficiently buffered, and capable of maintaining a physiologically neutral pH even when large amounts of lactate are produced (Robert A Robergs et al., 2004). Nonetheless, within poorly vascularised and fast-growing tumor masses, or in laboratory *ex vivo* cultures, lactate export results in acidosis, which has deleterious effects on T cell proliferation and survival (Brand et al., 2016; Fischer et al., 2007; Haas et al., 2015; Mendler et al., 2012). As a result, lactate has long been regarded as immunosuppressive, and many studies of lactate in immunity have focused on lactic acid, without considering the distinct role of the lactate anion itself. Thus, in order to discern the consequences of acidosis from those of increased lactate levels, we investigated the effect of a pH-neutral form of lactate, i.e., sodium lactate, during the initial phase of CD8+ T cell activation, when many dynamic changes to T cell gene expression and metabolism are occurring.

A growing body of literature shows that lactate has a wide range of functions in the organism. These include its roles as energy source, gluconeogenic precursor, signaling molecule, and epigenetic modifier (Zhang et al., 2019). For example, inter-cellular exchange of lactate occurs between white and red muscle fibers, skeletal muscle and heart, kidney and liver, as well as neurons and astrocytes (Baltazar et al., 2020; Brooks, 2020). In addition, recent studies have reported evidence of active lactate uptake as a carbon source in a wide variety of physiological and pathological circumstances (Chen et al., 2016; Faubert et al., 2017; Hui et al., 2018; Watson et al., 2021). Therefore, we wanted to characterise the fate and role of lactate in a metabolically dynamic, highly responsive system, i.e., CD8+ T cells.

CD8+ T cell metabolism and function are intimately connected (O’Neill et al., 2016; Pearce, 2010). Intracellular metabolism has been shown to drive and define T cell identity and immune activity (O’Neill et al., 2016). T cells are able to substantially rewire their energetic pathways and integrate different substrates in order to meet their specific energetic demands. For example, cytotoxic T cells may have to compete with cancer cells for nutrients within the tumor microenvironment (Chih-Hao Chang et al., 2015; Chih-Hao Chang and Erika L Pearce, 2016; Ping-Chih Ho et al., 2015). Hence, the ability to utilize alternative carbon sources can confer a proliferative advantage. In addition to their use of glucose as a primary energy source, CD8+ T cells heavily rely on glutamine and other amino acids such as leucine and serine to sustain effector functions (Carr et al., 2010; Hayashi et al., 2013; Ma et al., 2019; Manley et al., 2013).

We have previously shown that lactate is a critical factor in exercise-induced shifts in T cell surveillance of tumor growth (Rundqvist et al., 2020). Given the prevalence of lactate in the context of T cells undergoing priming, as well as in conditions of reduced oxygen availability in environments where T cells reside and traffic, we sought to determine the effect of the lactate anion, independently of pH, on CD8+ T cell metabolism and differentiation.

## Results

### Sodium lactate, but not lactic acid, is well-tolerated by CD8+ T cells

Upon TCR triggering, CD8+ T cells undergo a rapid induction of glycolytic metabolism which is followed by an increase in both intracellular and extracellular lactate levels (Supplementary Figure 1A-B). During glycolysis, protonated lactate, i.e., lactic acid, is released into the extracellular environment. The subsequent dissociation of the lactate anion from the proton leads to a localized decrease in pH (Figure 1A). In order to test the effect of lactate on CD8+ T cell activation and differentiation independently of pH reduction, we activated purified mouse CD8+ T cells *ex vivo* for 3 days in the presence of increasing doses of lactic acid, or its pH-neutral salt sodium lactate, using sodium chloride as a control for osmolality (Figure 1B and Supplementary Figure 1C). This allowed us to distinguish between the effects of the lactate anion itself and the acidification caused by proton dissociation. Addition of lactic acid to culture media caused a sharp decrease in pH, while equimolar doses of sodium lactate or sodium chloride did not alter media pH (Figure 1C). As reported previously (Brand et al., 2016), pH-lowering lactic acid has deleterious effects on CD8+ T cell activation, resulting in significantly reduced proliferation and cell numbers (Figure 1D and E). In contrast, activation in the presence of pH-neutral sodium lactate did not impair cell expansion, and had only a minimal effect on cell proliferation (Fig. 1D-F). A decrease in proliferation was observed at NaCl and sodium lactate concentrations above 50 mM, likely due to increased osmolality. Despite a slight delay in proliferation, exposure to 40 mM sodium lactate during the first 72 hours of activation was well tolerated by mouse CD8+ T cells (Figure 1F and Supplementary Figure 1D). Human CD8+ T cells activated in the presence of increasing doses of sodium lactate or sodium chloride showed a higher sensitivity to the rise in osmolality, but overall, a similar response to that seen in murine cells, indicating that lactate is also well tolerated by human CD8+ T cells (Supplementary Figure 1E and F).

**Figure 1.**
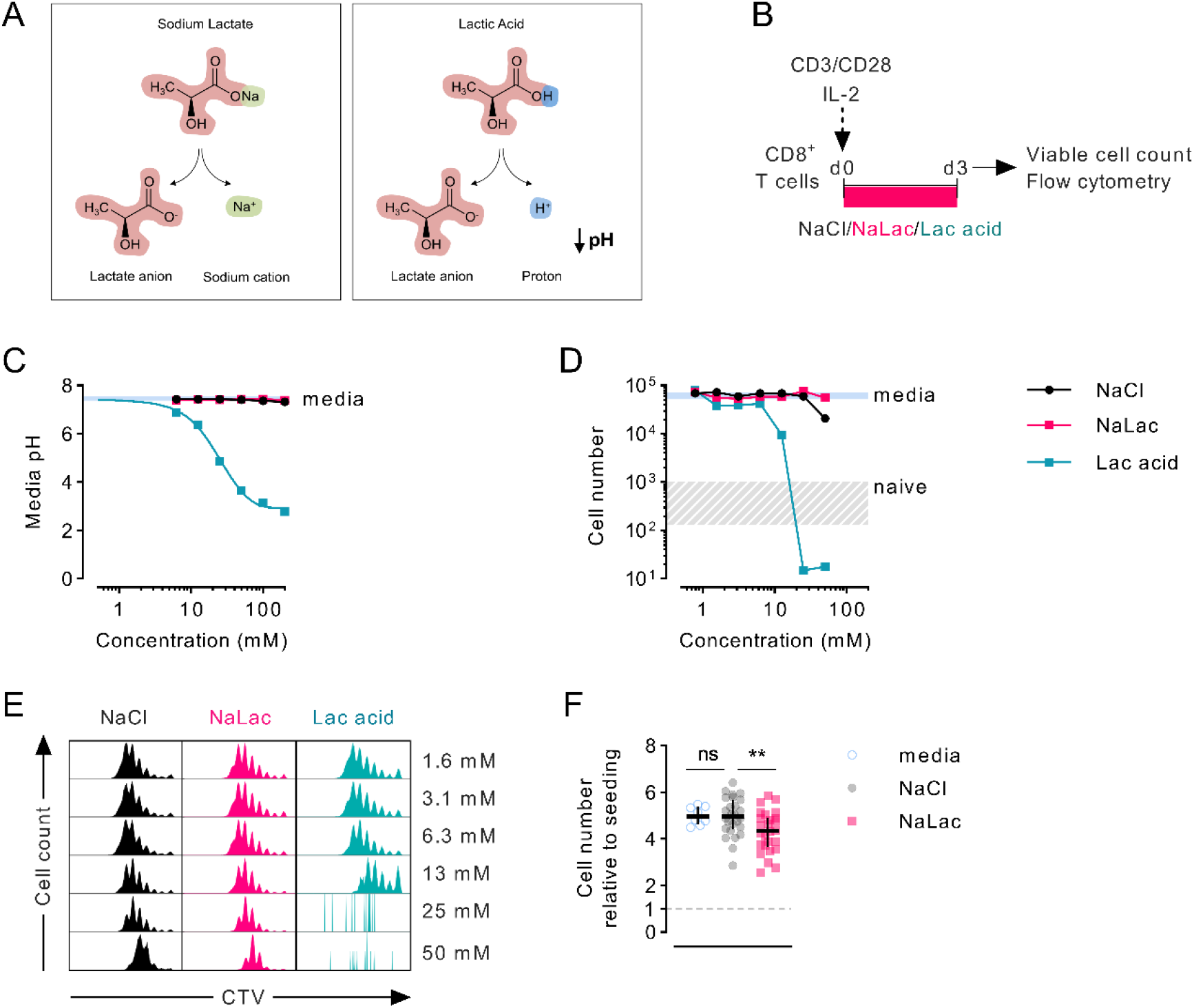
Sodium lactate, but not lactic acid, is well-tolerated by CD8+ T cells. **(A)** Diagram representing the dissociation of sodium lactate (NaLac, left) and lactic acid (Lac acid, right) in aqueous solution. **(B)** Experimental design. Mouse CD8+ T cells were activated for 72 hours in the presence of varying concentrations of sodium chloride (NaCl), sodium lactate (NaLac), or lactic acid (Lac acid). After activation, culture media was removed, cells were washed, counted and analysed by flow cytometry. **(C)** pH at room temperature of NaCl, NaLac, and Lac acid solutions in complete RPMI. Horizontal blue line: media reference. **(D)** Effect of increasing concentrations of NaCl, NaLac, and lactic acid on CD8+ T cell number, 72 hours after activation. Grey area: range of cell number for naive (non-activated) cells; Solid blue line: average of non-treated (media) CD8+ T cells. N = 1 composed of two mouse spleens pooled together. **(E)** Flow cytometry histograms from experiment in panel (C), showing cellular division profile by means of CellTrace Violet (CTV) dilution. **(F)** Number of mouse CD8+ T cells, 72 hours after activation in the presence of 40 mM NaCl, or NaLac, or media alone. Data are the median and interquartile range of n = 26 biological replicates. Cell number is presented relative to the number of cells seeded (dotted line). ** P<0.001 Dunnett’s multiple comparisons test.

### Lactate modulates CD8+ T cell metabolism

We next sought to characterize the immediate effects of lactate exposure on CD8+ T cell metabolism. To this end, we activated CD8+ T cells for 12 hours before carrying out real-time metabolic analyses, during which cells were exposed to 40 mM sodium lactate (Figure 2A). Relative to cells treated with media or an equimolar dose of NaCl, addition of sodium lactate reduced the extracellular acidification rate (ECAR) of activated CD8+ T cells and concomitantly increased the oxygen consumption rate (OCR), surrogate measurements for glycolysis and oxidative respiration, respectively (Figure 2B and C). Interestingly, despite suppressing glycolysis, addition of sodium lactate only slightly reduced the glycolytic capacity of CD8+ T cells (Figure 2B and C). This effect was also observed in human CD8+ T cell cultures (Supplementary Figure 2A-C). These results suggest that exogenous lactate is taken up by CD8+ T cells and induces metabolic alterations.

**Figure 2.**
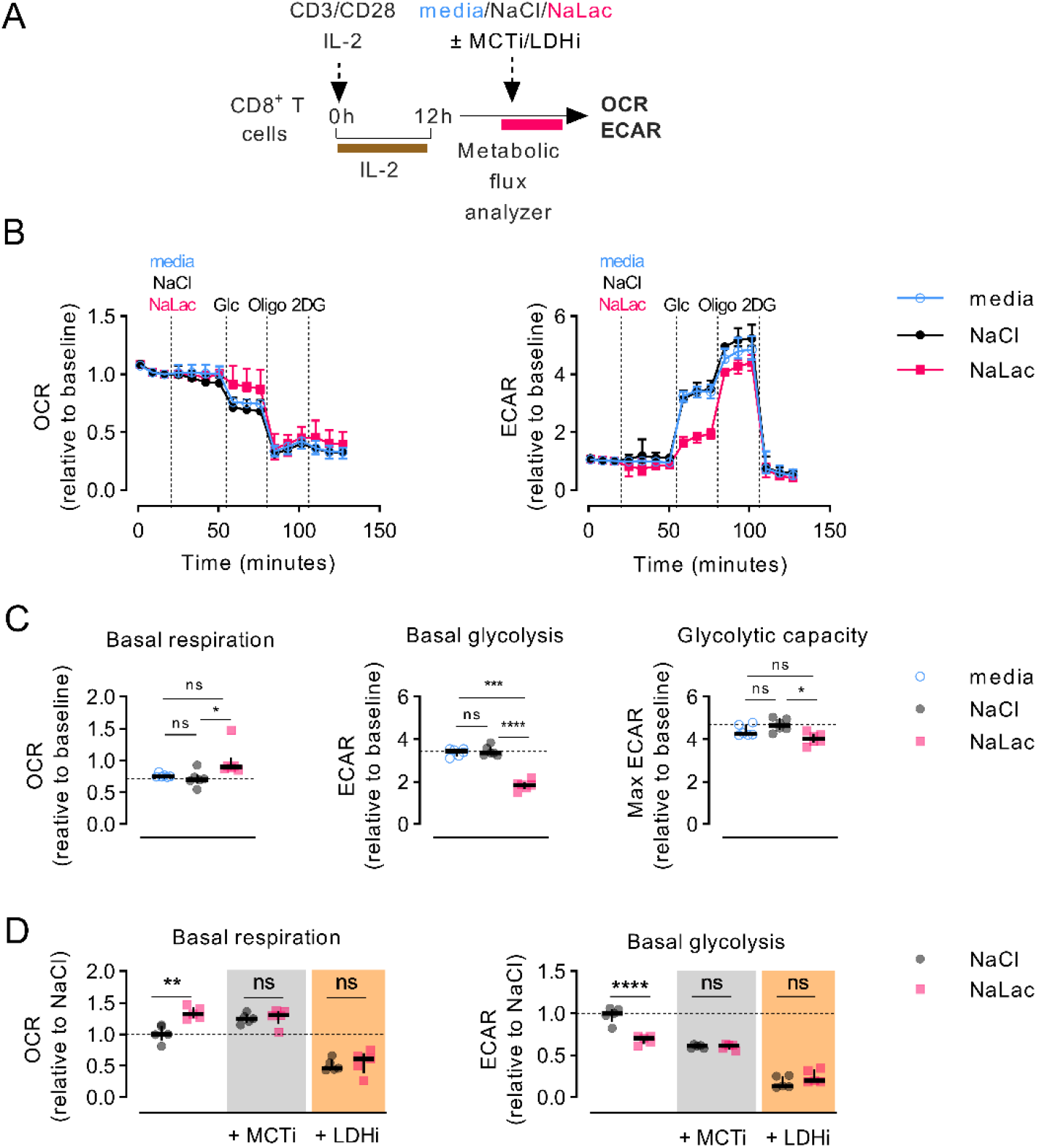
Lactate modulates CD8+ T cell metabolism. **(A)** Experimental design. CD8+ T cells were purified from mouse splenocytes and activated with anti-CD3/CD28 beads and IL-2 in media. After 12 hours, cells were counted, transferred into a metabolic flux analyzer, and exposed to either 40 mM NaCl, 40 mM sodium lactate (NaLac), or plain media prior to undergoing a glycolytic stress test. **(B)** Glycolytic stress test of mouse CD8+ T cells 12 hours after activation. Extracellular acidification rate (ECAR) and oxygen consumption rate (OCR) during injection of either media, 40 mM NaCl, or 40 mM NaLac, followed by 10 mM glucose, 1 μM oligomycin (oligo) and 50 mM 2-deoxyglucose (2-DG). ECAR and OCR are normalized to baseline levels. Data are the median and interquartile range of n = 6 independent mice. **(C)** Basal mitochondrial respiration, basal glycolysis and glycolytic capacity determined from the assay shown in (B). Dotted line: median of NaCl-treated cells. Data is the median and interquartile range of n = 6 independent mouse donors, each assayed as 4 technical replicates. * P<0.01, *** P<0.0005, **** p<0.0001 RM one-way ANOVA with Tukey’s multiple comparisons test. **(D)** Basal mitochondrial respiration and basal glycolysis determined from the represented shown in (A), after injection of either NaCl or NaLac with or without addition of 10 nM of the monocarboxylate transporter-1 (MCT-1) inhibitor AZD3985 or 50 μM of the lactate dehydrogenase (LDH) inhibitor GSK2837808A. Data are the median and interquartile range of n = 5 independent mouse donors. ** P<0.001, **** p<0.0001 two-way ANOVA with Šídák’s multiple comparisons test

The lactate anion is transported across cell membranes via the proton-linked monocarboxylate transporters (MCT-1, 2, 3, and 4) (Halestrap and Meredith, 2003; Halestrap and Price, 1999). Of these, only MCT1 and MCT4 are expressed in CD8+ T cells (Hukelmann et al., 2016; Tracy S P Heng et al., 2008; Wang and Green, 2012). While neither transporter is expressed at high levels in naïve CD8+ T cells, *ex vivo* activation causes MCT1 mRNA and protein to peak at 12 to 24 hours post-activation, whereas MCT4 expression peaks later, between 48 and 72 hours after activation (Supplementary Figure 2D and E). Activation of CD8+ T cells in the presence of an MCT1-selective inhibitor (AZD3965) inhibited cell proliferation at the low nanomolar range, irrespective of the presence of lactate (Supplementary Figure 2F), suggesting a crucial role for monocarboxylate transport in early T cell expansion. To confirm that the metabolic effect observed was due to uptake of exogenous lactate, we assessed the glycolytic and oxidative rates of T cells after administration of lactate in combination with the MCT1 inhibitor (Figure 2A). Blocking the molecular traffic of lactate via MCT1 mimicked the effects of sodium lactate, causing reduced ECAR and increased OCR (Figure 2D). This suggests that retention of endogenously generated lactate impacts T cell metabolism in a similar way to increases in exogenous lactate. As shown in figure 2D, adding exogenous sodium lactate during treatment with the MCT1 inhibitor did not further alter these T cell metabolic parameters.

The lactate dehydrogenase enzyme (LDH) plays a crucial role in lactate metabolism as it catalyzes the interconversion between pyruvate and lactate. Activated CD8+ T cells express high levels of LDHA and lower levels of LDHB (Supplementary Figure 2G) (Howden et al., 2019). We hypothesised that inhibition of LDH would prevent incorporation of exogenous lactate into carbon metabolism and abrogate its effects. However, inhibition of LDH activity with the LDH inhibitor GSK2837808A (Billiard et al., 2013) caused a drastic reduction of T cell expansion (Supplementary Figure 2H), and led to suppression of both basal glycolytic and oxidative respiration rates, in both the presence and absence of exogenous lactate (Figure 2D). This result highlights the central role played by LDH in the metabolism of activated CD8+ T cells, and indicates that disrupting the payoff phase of the glycolytic cascade has dramatic consequences on the metabolic fitness of the cell. However, inhibition of LDH did not allow us to determine if the effects of exogenous lactate on CD8+ T cell metabolism are directly dependent on the LDH-mediated conversion to pyruvate.

### Early exposure to lactate alters CD8+ T cell metabolism and gene expression

In order to better understand the role of lactate in CD8+ T cell metabolism, we carried out transcriptomic analysis of CD8+ T cells 72 hours after activation in the presence of 40 mM sodium lactate; these were coupled to metabolite quantification by mass spectrometry, as well as to metabolic flux analysis (Figure 3A). TCR triggering causes a drastic reprogramming of the cellular transcriptome, which, as shown, is further altered by the presence of lactate (Figure 3B and Supplementary Figure 3A). Lactate induced a concerted upregulation of genes involved in the TCA cycle. In contrast, expression of genes involved in glycolysis was not uniformly affected by lactate compared to media alone (Figure 3C and Supplementary Figure B).

**Figure 3.**
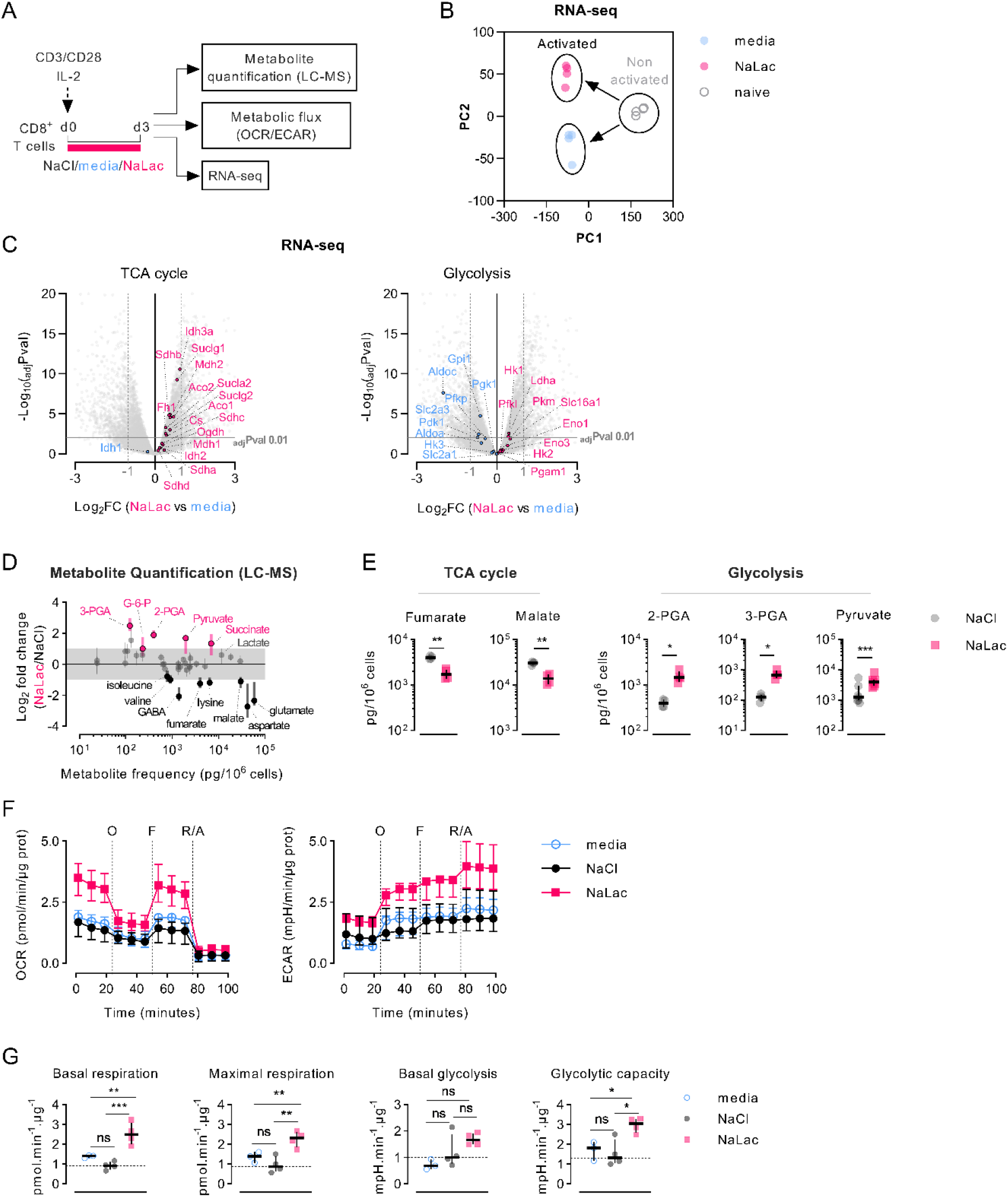
Early exposure to lactate alters CD8+ T cell metabolism and gene expression. **(A)** Experimental design. CD8+ T cells were purified from mouse splenocytes and activated with anti-CD3/CD28 beads and IL-2 in the presence of either plain media, or 40 mM NaCl, or 40 mM NaLac for 72 hours. Cells were then analyzed by liquid chromatography-mass spectrometry (LC-MS), metabolic flux analysis (OCR/ECAR), and RNA-sequencing. **(B)** Principal Component Analysis (PCA) plot of the RNA-seq data. Each dot represents a sample and each color represents a treatment. Light blue: mouse CD8+ T cells activated in plain media; pink: cells activated with 40 mM NaLac; grey: non-activated (naïve) cells. RNA was purified from 4 independent mouse donors per condition. Data is provided as Figure 3-source data 1. **(C)** Volcano plots generated from the RNA-seq data, representing differentially expressed TCA cycle (left) or glycolytic (right) genes in NaLac-relative to media-treated CD8+ T cells, 72 hours after activation. Significantly up- or down-regulated genes are highlighted respectively in pink or light blue. P values were calculated using Student’s t-test and adjusted by false discovery rate (FDR). Significance was set at adjusted p value (adjPval) ≤ 0.01, as indicated by the grey horizontal line. Dashed vertical lines indicate the threshold of differentially expressed transcripts (Log_2_ fold change ≥ 1 and ≤ −1). Data is provided as Figure 3-source data 2. **(D)** Metabolic changes after 72 hours-activation of mouse CD8+ T cells in the presence of NaLac compared to NaCl. Metabolites were extracted and quantified by Sugar Phosphate and Gas Chromatography-Mass Spectrometry (GC-MS). Pyruvate levels were determined by colorimetric assay. Data show log_2_ fold changes between NaLac- and NaCl-treated samples of n = 4 independent mouse donors. Pink and black dots represent significantly increased or decreased metabolite frequency in NaLac-versus NaCl-treated cells, respectively. Grey area: range between 2-fold increase and 2-fold decrease. Data is provided as Figure 3-source data 3. **(E)** Paired analysis of significant metabolites from the same assay as in (D), grouped by metabolic pathway (n = 4, with each pair representing an independent mouse donor). Pyruvate levels were determined by a colorimetric assay from n = 7 independent mouse donors. ** P<0.01, * P<0.05 twotailed paired t test. **(F)** Oxygen consumption rate (OCR) and extracellular acidification rate (ECAR) of CD8+ T cells after 72 hours activation in the presence of 40 mM NaCl, 40 mM NaLac, or plain media. OCR and ECAR were measured during sequential addition of 1 μM oligomycin (oligo), 1.5 μM FCCP, and 100 nM rotenone combined with 1 μM antimycin A (Rot/Ant). OCR and ECAR were normalized to total protein content at the end of assay. Lines represent the median and interquartile range of n = 4 independent mouse donors, each assayed as 4 technical replicates. **(G)** Basal respiration, maximal respiration, basal glycolysis, and glycolytic capacity determined from the assay shown in (F). Dotted line: median of NaCl-treated (control) cells. * P<0.05, ** P<0.001, *** P<0.001 one-way ANOVA with Tukey’s multiple comparison test.

Mass spectrometric analysis of the T cell metabolome revealed increased levels of glycolytic intermediates, including glucose-6-phosphate (G-6-P), 3- and 2-phosphoglyceric acid (3-PGA, 2-PGA), and pyruvate, in CD8+ T cells exposed to sodium lactate for 3 days (Figure 3D, E and Supplementary Figure 3C). Within the TCA cycle intermediates, lactate exposure caused an increase in the amount of succinate as well as a reduction in the downstream metabolites fumarate and malate (Figure 3D, E and Supplementary Figure 3C). Decreased levels of anaplerotic metabolites (glutamate and aspartate) were also observed. Mitochondrial stress tests confirmed that CD8+ T cells activated in the presence of lactate for 72 hours exhibit higher basal and maximal respiration as well as increased glycolytic rates and glycolytic capacity at the end of the exposure (Figures 3F and G). Altogether, these results demonstrate that CD8+ T cells adapt transcriptionally and metabolically to exposure to lactate during activation.

### Lactate is incorporated in cellular metabolism and displaces glucose

To determine if sodium lactate is taken up by CD8+ T cells and incorporated into carbon metabolism, we activated CD8+ T cells in the presence of 11.1 mM [U-^13^C_6_]glucose alone, or in combination with 40 mM sodium lactate, or 40 mM [U-^13^C_3_]sodium lactate in combination with 11.1 mM glucose (Figure 4A and Supplementary Figure 4). Labelling of cellular metabolites was measured by mass spectrometry at 24, 48 and 72 hours after activation (Figure 4B-D and Supplementary Figure 4). Activation in the presence of [U-^13^C_6_]glucose alone resulted initially (24-48h) in labelling of metabolites in the glycolytic pathway, including phosphoenolpyruvate (PEP), then pyruvate and lactate, followed by TCA cycle intermediates at 48-72 hours. Progressive labelling of ribose, originating from the pentose phosphate pathway (PPP), and the amino acids serine, glycine, alanine, glutamate, proline, and aspartate was also observed, suggesting incorporation of glucose-derived carbons through glycolysis and its branch pathways, as well as the TCA cycle. Addition of sodium lactate significantly displaced the contribution of [U-^13^C_6_]glucose to the labelling of pyruvate and lactate, but not to PEP or ribose (Figure 4C and Supplementary Figure 4). Glucose labelling of TCA cycle metabolites was markedly decreased by addition of lactate, as was the labelling of amino acids (Figure 4C and Supplementary Figure 4). Tracing of [U-^13^C_3_]sodium lactate revealed that exogenously added lactate enters T cells and is oxidized to pyruvate (Figure 4D and Supplementary Figure 4). Labelling of metabolites downstream of pyruvate, namely alanine and citrate, occurs rapidly. Within 24 hours, labelling from lactate-derived carbons can be found in the TCA cycle metabolites succinate, fumarate and malate. In addition to alanine, lactate-derived carbons were incorporated into amino acids derived from the TCA cycle such as glutamate, proline and aspartate, but not into amino acids derived from the glycolytic pathway upstream of pyruvate, e.g., serine and glycine (Figure 4D and Supplementary Figure 4). These findings show that exogenous lactate can be taken up, oxidized, and displace glucose as a carbon source for both the TCA cycle and amino acid synthesis in activated CD8+ T cells.

**Figure 4.**
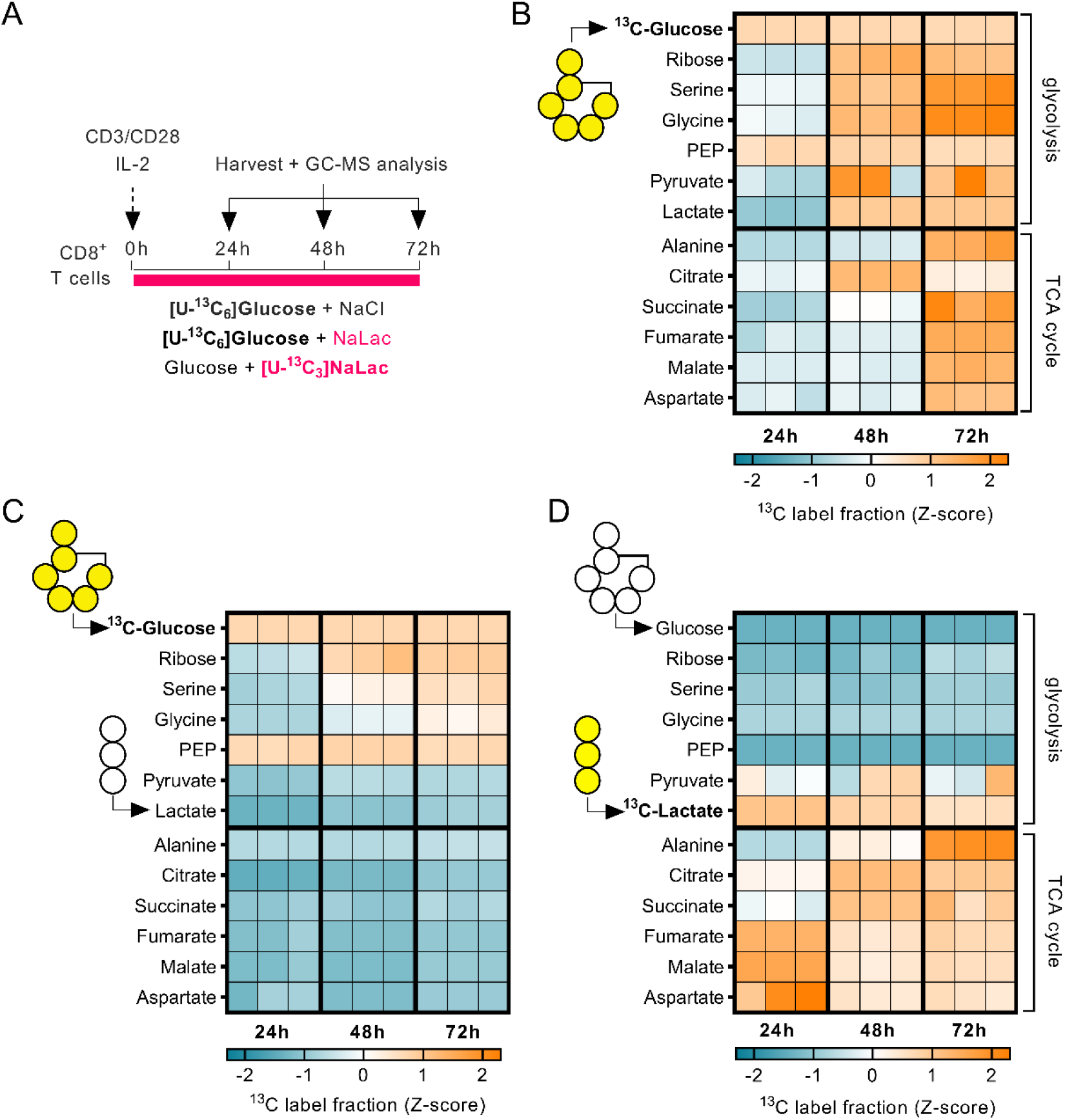
Lactate is incorporated in cellular metabolism and displaces glucose. **(A)** Experimental design. Mouse CD8+ T cells were activated in the presence of either 11.1 mM [U-^13^C_6_]glucose + 40 mM NaCl, or 11.1 mM [U-^13^C_6_]glucose + 40 mM sodium lactate, or 11.1 mM glucose and 40 mM [U-^13^C_6_]sodium lactate. Incorporation of labelled carbons in cellular metabolites was assessed by GC-MS at 24, 48, and 72 hours after activation. **(B-D)** Heatmaps showing glucose- (B, C) or lactate-derived (D) carbons incorporation into cellular metabolites at 24, 48, and 72 hours after activation, expressed as Z-scores. N = 3 individual mouse donors, each one represented by a column at each time-point. Data is provided as Figure 4-source data 1.

### Activation in the presence of lactate modulates the differentiation of CD8+ T cells

In addition to metabolic genes, analysis of the cell transcriptome after exposure to lactate revealed a substantial effect on the expression of genes involved in CD8+ T cell differentiation and cytotoxic activity, including Granzyme B (Gzmb), interferon-gamma (IFN-γ, IFNg), perforin 1 (Prf1) and ICOS (Icos) (Figure 5A). Indeed, addition of sodium lactate during activation elicited a dose-dependent increase in GZMB expression, beginning at 10 mM and peaking at 40-50 mM (Figure 5B). Granzyme B is one of the most highly expressed effector proteins in activated mouse CD8+ T cells (Hukelmann et al., 2016) and is essential to granule-mediated apoptosis of target cells (Jonathan W. Heusel et al., 1994). In addition to a significant increase in GZMB expression (Figure 5C and Supplementary Figure 5A and B), mouse CD8+ T cells activated with 40 mM sodium lactate showed increased secretion of interferon-gamma, another important cytotoxic mediator (Figure 5D). Lactate also altered expression of a number of other proteins involved in CD8+ T cell differentiation, increasing the levels of CD44, CTLA-4, 4-1BB, and ICOS, and lowering the levels of the transcription factor T-bet and the lymph node homing molecule CD62L (Figure 5D and Supplementary Figure 5C).

**Figure 5.**
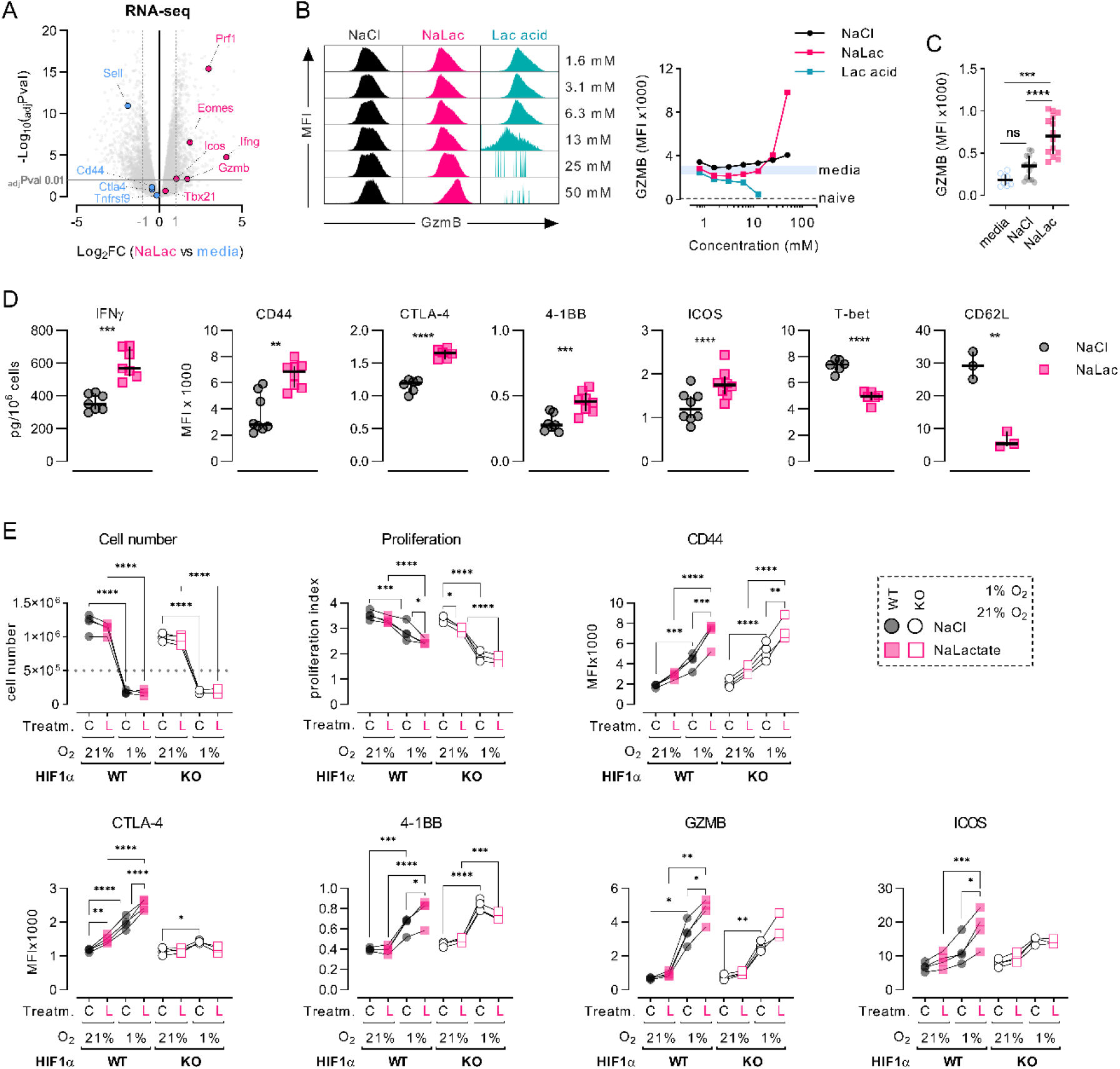
Activation in the presence of lactate modulates the differentiation of CD8+ T cells. **(A)** Volcano plot of the RNA-seq data, showing genes associated with T cell differentiation and function that are differentially expressed in NaLac-versus media-treated, activated CD8+ T cells. Significantly up- or down-regulated genes (Log_2_ fold change ≥ 1 or ≤ −1) are highlighted in pink or light blue, respectively. P values are calculated from Student’s t-test and adjusted by false discovery rate (FDR). Significance is set at adjusted p value (adjPval) ≤ 0.01. Grey horizontal line indicates adjPval = 0.01. Dashed vertical lines indicate the threshold of differentially expressed transcripts (Log_2_ fold change ≥ 1 and ≤ −1). Data is provided as Figure 5-source data 1. **(B)** CD8+ T cells were activated for 72 hours in the presence of increasing doses of sodium chloride (NaCl), sodium lactate (NaLac), or lactic acid (Lac acid). After activation, intracellular granzyme B (GZMB) levels were analysed by flow cytometry. Left: histograms showing the Median Fluorescence Intensity (MFI) of GZMB. Right: MFI of GZMB at increasing doses of the different treatments. Solid light blue line: range (minimum to maximum) of media-treated, activated CD8+ T cells (control); dotted line: average of non-activated CD8+ T cells (naïve). **(C)** Median Fluorescence Intensity (MFI) of intracellular Granzyme B (GZMB) measured by flow cytometry, at day 3 after activation with 40 mM NaCl, 40 mM of NaLac, or plain media for 72 hours. Data are the median and interquartile range of n = 17 biological replicates. **** P<0.0001 Dunnett’s multiple comparisons test. **(D)** Levels of secreted interferon-γ (IFN-γ) and Median Fluorescence Intensity (MFI) of CD44, CTLA-4, 4-1BB, ICOS, T-bet and CD62L in mouse CD8+ T cells activated for 72h in the presence of 40 mM NaCl or 40 mM NaLac. Each pair represents an independent mouse donor (n = 3-9). ** P<0.01, *** P<0.001, **** p<0.0001 two-tailed paired t-test. **(E)** Cell number, Division Index, Proliferation Index, and Median Fluorescence Intensity (MFI) of surface markers CD44, CTLA-4, 4-1BB, and ICOS, and intracellular GZMB levels in wild type (Hif1 α^fl/fl^) or HIF-1α deficient CD8+ T cells (Hif1α^fl/fl^ dLck Cre) activated for 3 days in the presence of either 40 mM NaCl (C) or NaLac (L), at 1% or 21% O_2_. Grey dotted line: number of seeded cells. N = 4 independent mouse donors. * p<0.05, ** p<0.005, *** p<0.0005, ordinary one-way ANOVA with Šídák’s multiple comparisons test. Non-significant comparisons not indicated.

Because lactate production is increased in hypoxic conditions and since this is in part driven by the Hypoxia Inducible Factor-1α (HIF-1α) transcription factor, we measured the differentiation of CD8+ T cells activated in the presence of 40 mM sodium lactate at ambient oxygen tension (21% O_2_) or hypoxic conditions (1% O_2_) using wild type or HIF-1α null CD8+ T cells (Figure 5E). Activation in hypoxic conditions severely limited CD8+ T cell proliferation but led to greater expression of the effector markers CD44, CTLA-4, 4-1BB, GZMB and ICOS, in agreement with previously published data (Doedens et al., 2013; Palazon et al., 2017). Exposure to lactate led to a further increase in expression of these markers; this was enhanced in hypoxia. Effector differentiation has been shown to be compromised in the absence of HIF-1α (Doedens et al., 2013; Finlay et al., 2012; Palazon et al., 2017). We observed that CD8+ T cells lacking HIF-1α failed to upregulate CTLA-4 in any of the tested culture conditions, suggesting a high level of HIF-1α-dependency. In contrast, expression of CD44, 4-1BB, GZMB and ICOS was increased in hypoxia even in the absence of HIF-1α. The additive effect of lactate and hypoxia on the expression of 4-1BB and ICOS was lost in HIF-1α null CD8+ T cells; this HIF-dependency was not seen in GZMB and CD44. Altogether, these results show that lactate and hypoxia have additive effects in driving expression of CD8+ T cell effector markers but indicate that that relationship may only in part be dependent on HIF-1α.

## Discussion

Here we show that the lactate anion is a non-toxic molecule that can be taken up and incorporated into T cell metabolism, and that by shaping the transcriptome, lactate can ultimately alter the fate of CD8+ T cells.

A wide range of mammalian tissues can utilize lactate as a carbon source to fuel their metabolism. In fact, in many cases, lactate is used as a preferred energy source (Brooks, 2018; Faubert et al., 2017; Hui et al., 2018). Rapidly proliferating cells, including malignant cells and activated lymphocytes as well as metabolically active tissues such as exercising muscle, break down large amounts of glucose in order to feed biosynthetic pathways and provide oxidative substrates to the TCA cycle. It is also clear that the generation of lactate from pyruvate is essential for continued glycolytic flow in order to regenerate the NAD+ cofactor (Heiden et al., 2009). Immune cells are often exposed to high levels of lactate, in both loci of disease and in metabolically active tissues, and this can also be associated with exposure to a localized reduction in tissue pH. Although healthy tissue microenvironments are capable of maintaining a physiologically neutral pH even when large amounts of lactate are produced (Péronnet and Aguilaniu, 2006), in poorly vascularized and fast-growing tumor masses, or in laboratory *ex vivo* cultures, lactate export can result in significant acidosis. It has been extensively demonstrated that tumor-derived lactate can inhibit the functions of T and NK cells due to this intracellular acidification (Brand et al., 2016). Likewise, lactate-associated acidity has been shown to impair CD8+ T cell motility (Haas et al., 2015). However, these studies do not distinguish between the effect of the lactate anion itself, and the pH-lowering properties of its protonated form, i.e., lactic acid.

Activating CD8+ T cells with carbon-labeled lactate allowed us to trace its active uptake and incorporation into carbon metabolism. After being oxidized to pyruvate, exogenous lactate was not incorporated into upstream metabolites in the glycolytic cascade. Instead, lactate-derived carbons were specifically incorporated into TCA cycle intermediates and pyruvate-derived amino acids such as alanine. Interestingly, this resulted in the displacement of glucose as the preferred source of carbons for TCA intermediates. Thus, our data indicate that exogenous lactate, at least when present in high doses, replaces glucose as the preferred substrate to generate specific amino acids and TCA cycle intermediates. Unexpectedly, the presence of lactate seems to indirectly impair how glucose feeds the serine, glycine, one-carbon (SGOC) metabolism, even though no contribution of lactate-derived carbons could be detected in either glycine or serine. Recently, this metabolic network has been linked to CD8+ T cells effector functions (Ma et al., 2017). Hence, by altering the intracellular flow of glucose, lactate might modulate the overall immunological activity of CD8+ T cells.

Increased glucose uptake and processing through the glycolytic pathway results in a sharp increase in lactate production. Elevated levels of lactate in the microenvironment might act as a feedback mechanism to reprogram T cell metabolism (Zhang et al., 2019). Our data indicate that lactate accumulation represses glycolytic flux whilst boosting mitochondrial respiration.

Lactate enhanced the expression of cytotoxic molecules such as Granzyme B and Interferon-γ. This was paired with a reduction in surface CD62L, a lymph node homing marker, and enhanced expression of activation markers including surface molecule CD44, co-activators 4-1BB and ICOS, and co-inhibitor CTLA-4. Lactate also reduced the levels of a master regulator of CD8+ T cell differentiation, the transcription factor T-bet. Previous work has shown that HIF-1α is necessary for effector differentiation of CD8+ T cells (Palazon et al., 2017). Our data indicate that lactate and hypoxia have additive effects in shaping CD8+ T cell fate by a mechanism that is only partially dependent on HIF-1α.

Overall, our work reveals a possible mechanism of how lactate is utilised within CD8+ T cells. We show that exogenously supplied lactate is imported and used as a carbon source to fuel intracellular metabolism. The use of distally produced endogenous lactate as a fuel for oxidative respiration has been previously reported in other cell types, both malignant and non-malignant. Our data suggest that microenvironmental lactate can also be used by CD8+ T cells, at least during the early effector differentiation phase. Locally-produced lactate originating from primed T cells might, for example, function as a quorum-sensing mechanism to cross-potentiate the early T cell response. The exceptional plasticity of T cells represents one of their most distinctive features, and the ability to rapidly adapt to diverse environmental conditions, including reduced oxygen availability and nutrient deprivation, is of crucial importance for their immune activity. As we have recently shown that administration of sodium lactate to mice inhibits tumor growth in a T cell-dependent manner (Rundqvist et al., 2020), our findings here indicate that lactate has a potential function as an immunomodulatory metabolite in immunotherapy.

## Material and Methods

### Animals

All experiments and protocols were approved by the regional animal ethics committee of Northern Stockholm (dnr N78/15, N101/16). For T cell purification, wild type donor and recipient C57BL/6J animals were purchased from Janvier Labs.

### Osmolality measurements

Sodium Chloride (Sigma, S5886) and Sodium L-Lactate (Sigma, L7022) solutions were prepared by dissolving each compound into RPMI 1640 (Thermo Fisher, 21875) at a maximum concentration of 500 mM. Solutions were serially diluted and osmolality measurements were obtained with a micro-osmometer (Löser Messtechnik, D-1000 Berlin 38).

### Mouse CD8+ T cell purification, activation and expansion

Spleens were harvested from 8-12 week old C57BL/6J mice, mashed over a 40 μm cell strainer (VWR, 10199-654), and CD8+ T cells were purified by positive magnetic bead selection (Miltenyi Biotec, 130-117-044) according to manufacturer’s instructions. Purified CD8+ T cells were counted and cell diameter measured using a Moxi Z mini (Orflo, MXZ001) or a TC20 (Bio-Rad, 145-0101) automated counter. 1×10^6^ (24-well plate) or 5×10^5^ (48-well plate) CD8+ T cells were activated with Dynabeads Mouse T-Activator CD3/CD28 (Thermo Fisher, 11456D) in a 1:1 bead to T cell ratio and cultured in 2 ml (24-well plate) or 1 ml (48-well plate) RPMI 1640 (Thermo Fisher, 21875) supplemented with 10% Fetal Bovine Serum (Thermo Fisher, 10270-106), 50 μM 2-mercaptoethanol (Thermo Fisher, 21985023), 100 U/ml penicillin-streptomycin (Thermo Fisher, 15140122), and 10 U/ml recombinant human IL-2 (Sigma, 11011456001), and incubated at 37° C for 3 days in a humidified CO_2_ incubator.

### Human CD8+ T cell purification, activation and expansion

Peripheral blood mononuclear cells (PBMCs) were harvested from standard buffy coat preparations of healthy donors, aged 20 to 40 years old, obtained from the Department of Transfusion Medicine at Karolinska University Hospital and processed no later than 4 hours after collection. PBMCs were isolated by gradient centrifugation using Histopaque-1077 (Sigma, 10771) and enriched for CD8+ T cells by magnetic bead selection, according to manufacturer’s instructions (Miltenyi Biotec, 130-045-201). Purified CD8+ T cells were counted and 5×10^5^ (48-well plate) or 1×10^5^ (96-well plate) naïve CD8+ T cells were activated with Dynabeads Human T-Activator CD3/CD28 (Thermo Fisher, 11132D) in a 1:1 bead to T cell ratio and cultured in 1 ml (48-well plate) or 200μl (96-well plate) RPMI 1640 (Thermo Fisher, 21875) supplemented with 10% Fetal Bovine Serum, (Thermo Fisher, 10270-106), 100 U/ml penicillin-streptomycin (Thermo Fisher, 15140122), and 30 U/ml recombinant human IL-2 (Sigma, 11011456001), and incubated at 37°C for 4 days in a humidified CO_2_ incubator. For long-term expansion after 4 days, Dynabeads were removed using a DynaMag-2 Magnet (Thermo Fisher, 12321D), washed with PBS, counted, and resuspended in fresh media supplemented with 30 U/ml IL-2 and incubated as described above. Media supplemented with cytokines was added every 2-3 days in order to maintain cell densities below 2×10^6^/ml. Cells were counted again at day 7 and maintained in culture under the above described conditions until day 14.

### CD8+ T cell *ex vivo* activation

Sodium L-Lactate (Sigma, L7022), L-Lactic Acid (Sigma, L1750) and Sodium Chloride (Sigma, S5886) were prepared as 10x concentrated solutions in complete media. Compounds were added to T cells at the point of activation (day 0) minutes before addition of CD3/CD28 Dynabeads. Sodium Chloride and/or plain media was used as control. For long-term expansion, after beads removal cells were cultured in complete media supplemented with cytokines without further addition of compounds.

### Cell proliferation analysis

After magnetic bead purification, CD8+ T cells were loaded with 5 μM CellTrace CFSE or CellTrace Violet (CTV) dyes (Thermo Fisher, C34554 and C34557, respectively) according to manufacturer’s instructions. CFSE or CTV dilution was determined by flow cytometry.

### Cell phenotyping by flow cytometry

Single-cell suspensions were washed and stained with Fixable Near-IR Dead Cell Stain Kit (Thermo Fisher, L10119) followed by staining of extracellular antigens with fluorochrome labelled antibodies. The Fixation/Permeabilization Solution Kit (BD Biosciences, 554714) was used for exposing cytoplasmic antigens. The Transcription Factor Buffer Set (BD Biosciences, 562574) was used for exposing nuclear antigens. Fluorochrome-labelled antibodies against mouse antigens CD44 (clone IM7), CD8 (clone 53-6.7), CTLA-4 (clone UC10-4F10-11), LAG-3 (clone C9B7W), and PD-1 (clone J43) were purchased from BD Biosciences, CD27 (clone LG.3A10), CD28 (clone 37.51), CD62L (clone MEL-14), and ICOS (clone C398.4A) were purchased from BioLegend, and 4-1BB (clone 17B5), CD127 (clone A7R34), CD25 (clone PC61.5), Eomes (clone Dan11mag), T-bet (clone eBio4B10), and anti-human/mouse Granzyme B (clone GB12) was purchased from Thermo Fisher. Cell counting was performed with CountBright Absolute Counting Beads (Thermo Fisher, C36950). Samples were processed in a FACS Canto II flow cytometer (BD Biosciences). Data analysis was performed with FlowJo, version 10.7.2.

### Western blotting and real time RT-PCR

Mouse CD8+ T cells were activated *ex vivo* and cells harvested at 0, 6, 12, 24, 48 and 72 hours post-activation. For each time point, total protein was extracted using the NE-PER™ Nuclear and Cytoplasmic Extraction Reagents kit (78833, Thermo Fisher) and the cytoplasmic fraction used for western blots. Total RNA was extracted using the RNeasy Mini Kit (74104, Qiagen). For each sample, 15 μg of total protein were separated in SDS-PAGE, blotted onto a PVDF membrane and probed with antibodies against MCT1 (sc-50325, Santa Cruz), MCT4 (sc-50329, Santa Cruz), GzmB (4275, Cell Signaling Technologies) and PPIB (A7713, Antibodies Online) and detected using infra-red labeled secondary antibodies in an Odyssey imaging system (LICOR). REVERT Total protein stain (926-11010, Li-COR) was used for lane normalization. Two micrograms of total RNA were reverse transcribed using iScript cDNA synthesis kit (BioRad) in a total volume of 20 μl. Real-time RT-PCR was used for mRNA quantification (7500 Fast Real-Time PCR system, Applied Biosystems Inc., Foster City, California, USA). All primers were designed to cover exon-exon boundaries to avoid amplification of genomic DNA. Predesigned qPCR primers used to detect mouse Slc16a1 (forward: 5’-ACTTGCCAATCATAGTCAGAGC-3’, reverse: 5’-CGCAGCTTCTTTCTGTAACAC-3’), Slc16a3 (forward: 5’-GACGCTTGTTGAAGTATCGATTG-3’, reverse: 5’-GCATTATCCAGATCTACCTCACC-3’), granzyme B (GzmB) (forward: 5’-CTGCTAAAGCTGAAGAGTAAG-3’, reverse: 5’-TAGCGTGTTTGAGTATTTGC-3’), and Hprt (forward: 5’-TGACACTGGCAAAACAATGCA-3’, reverse: 5’-GGTCCTTTTCACCAGCAAGCT-3’) were purchased from IdtDNA or Sigma. All reactions were performed in 96-well MicroAmp Optical plates in duplicates. The total reaction volume was 15 μl, containing 5 μl sample cDNA, 0.4 mM of each primer forward and SYBR Green PCR Master Mix (4309155E, Applied Biosystems Ins.). All quantification reactions were controlled with a melting curve and primer efficiency was tested with standard curves, and did not differ between the primer pairs. Gene expression data was normalized to Hprt levels.

### RNA sequencing

Mouse CD8+ T cells were activated as previously described, for 72 hours in the presence of either 40 mM sodium lactate or plain media. At the end of activation cells were washed twice with PBS and cell pellets were snap-frozen in RLT Plus lysis buffer (Qiagen, #1053393). Total RNA was extracted with the Qiagen’s RNeasy kit according to the manufacturer’s instructions. All samples were quality checked with Agilent Tapestation RNA screen tape. To construct libraries suitable for Illumina sequencing, the Illumina TruSeq Stranded mRNA Sample preparation protocol which includes cDNA synthesis, ligation of adapters, and amplification of indexed libraries was used. The yield and quality of the amplified libraries were analysed using Qubit by Thermo Fisher and the Agilent Tapestation. The indexed cDNA libraries were normalised and combined, and the pools were sequenced on the Nextseq 550 for a 50-cycle v2.5 sequencing run, generating 71 bp single-end reads (and 2*10 bp index reads). Demultiplexing was performed using bcl2fastq (v2.20.0.422), generating fastq files for further downstream mapping and analysis.

### RNA sequencing analysis

Bcl files were converted and demultiplexed to fastq using the bcl2fastq v2.20.0.422 program. STAR 2.7.5b (Dobin et al., 2013) was used to map the fastq files to the mouse reference genome (mm10/GRCm38) and to remove PCR duplicates. Uniquely mapped reads were then counted in annotated exons using featureCounts v1.5.1 (Liu et al., 2015). The gene annotations (Mus_musculus.GRCm38.99.gtf) and reference genome were obtained from Ensembl. The count table from featureCounts was imported into R/Bioconductor and differential gene expression was performed using the EdgeR package (Robinson et al., 2010) and its general linear models pipeline. For the gene expression analysis, genes that had 1 count per million in 3 or more samples were used and normalized using TMM normalization. Genes with an FDR adjusted p value ≤ 0.01 were termed significantly regulated. Principal Component Analysis (PCA) plots were obtained by normalizing gene counts after filtering out lowly expressed genes.

### Metabolite extraction

For metabolite analysis, 5×10^6^ cells were harvested and washed 3 times with PBS before adding 800 μL of ice-cold extraction mixture consisting of chloroform:MeOH:H2O (1:3:1), containing 500 pg/μL of 2-Deoxy-D-glucose 6-phosphate and 70 pg/μL of myristic acid-^13^C_3_ were added to each sample. Metabolites were extracted using a mixer mill set to a frequency 30 Hz for 2 min, with 1 tungsten carbide bead added to each tube. Obtained extracts were centrifuged at 14000 rpm for 10 min. The collected supernatants were divided for GC-MS and Sugar-P analysis. 300 μL of the supernatant was transferred into GC and LC-vial respectively, the supernatants were evaporated until dryness using a SpeedVac.

### Sugar phosphate analysis

For derivatization, dried samples were dissolved in 20 μl of methoxylamine and incubated on a heat block at 60°C, 30 min. After overnight incubation at room temperature, 12μl of 1-Methylimidazol and 6 μl of propionic acid anhydride were added and heated at 37°C for 30 minutes. The reaction mixture was then evaporated to dryness by N2 gas. Prior to LC-MS analysis, derivatized metabolites were dissolved in 100μl of aqueous 0.1% formic acid. Quantitative analysis was done by combined ultra-high-performance liquid chromatography electrospray ionization-triple quadrupole-tandem mass spectrometry (UHPLC-ESI-QqQMS/MS) in dynamic multiple-reaction-monitoring (MRM) mode. An Agilent 6495 UHPLC chromatograph equipped with a Waters Acquity BEH 1.7 μm, 2.1 x 100 mm column (Waters Corporation, Milford, USA) coupled to a QqQ-MS/MS (Agilent Technologies, Atlanta, GA, USA) was used. The washing solution, for the autosampler syringe and injection needle, was 90% MeOH with 1% HCOOH. The mobile phase consisted of A, 2% HCOOH and B, MeOH with 2% HCOOH. The gradient was 0% B for 1 min followed by linear gradients from 0.1 to 30% from 1 to 3 min then 30 to 40% B from 3 to 6 min, hold at 40% B from 6 to 10 min, followed by 40 to 70% B from 10 to 12.5 min, hold at 70% B from 12.5 to 15 min, and thereafter 70 to 99% B from 15 to 17.5 min. B was held at 99% for 0.5 min, and thereafter the column was re-equilibrated to 0% B. The flow rate was 0.65 mL min-1 during equilibration and 0.5 mL min-1 during the chromatographic runs. The column was heated to 40 °C, and injection volumes were 1 μL. The mass spectrometer was operated in negative ESI mode with gas temperature 230°C; gas flow 12 L min-1; nebulizer pressure 20 psi; sheath gas temperature 400°C; sheath gas flow 12 L min-1; capillary voltage 4000 V (neg); nozzle voltage 500 V; iFunnel high pressure RF 150 V; iFunnel low pressure RF 60 V. The fragmentor voltage 380 V and cell acceleration voltage 5 V. For a list of MRM transitions see Suppl. Data Table S1. Data were processed using MassHunter Qualitative Analysis and Quantitative Analysis (QqQ; Agilent Technologies, Atlanta, GA, USA) and Excel (Microsoft, Redmond, Washington, USA) software.

### GC-MS analysis

The GC-MS samples were spiked with 1050 pg of each GC-MS internal standard before evaporation. Derivatization was performed according to Gullberg *et al.* (Gullberg et al., 2004). In detail, 10μL of methoxyamine (15 μg/μL in pyridine) was added to the dry sample that was shaken vigorously for 10 minutes before being left to react at room temperature. After 16 hours 10 μL of MSTFA was added, the sample was shaken and left to react for 1 hour at room temperature. 10 μL of methyl stearate (1050 pg/μL in heptane) was added before analysis. One μL of the derivatized sample was injected by an Agilent 7693 autosampler, in splitless mode into an Agilent 7890A gas chromatograph equipped with a multimode inlet (MMI) and 10 m x 0.18 mm fused silica capillary column with a chemically bonded 0.18 μm DB 5-MS UI stationary phase (J&W Scientific). The injector temperature was 260 °C. The carrier gas flow rate through the column was 1 ml min-1, the column temperature was held at 70 °C for 2 minutes, then increased by 40 °C min-1 to 320 °C and held there for 2 min. The column effluent 16 is introduced into the electron impact (EI) ion source of an Agilent 7000C QQQ mass spectrometer. The thermal AUX 2 (transfer line) and the ion source temperatures were 250 °C and 230 °C, respectively. Ions were generated by a 70 eV electron beam at an emission current of 35 μA and analyzed in dMRM-mode. The solvent delay was set to 2 minutes. For a list of MRM transitions see Suppl. Data Table S2. Data were processed using MassHunter Qualitative Analysis and Quantitative Analysis (QqQ; Agilent Technologies, Atlanta, GA, USA) and Excel (Microsoft, Redmond, Washington, USA) software.

### Isotopic labelling, metabolite extraction and GC-MS analysis

Mouse CD8+ T cells were activated with CD3/CD28 beads and 10 U/ml IL-2 in glucose-free RPMI supplemented with 2 mM glutamine, 10% FBS, 100 U/ml penicillin/streptomycin and 2-mercaptoethanol, and either i) 11 mM [U^13^C_6_] glucose, ii) 11mM [U^13^C_6_] glucose and 40 mM lactate, or iii) 11 mM glucose and 40 mM [U^13^C_3_] sodium lactate, and isotopic labelling was performed for 24, 48 and 72 hours. Sodium chloride (40 mM) was added to i) to control for osmolality. Cells harvested at each time point were washed with PBS, and metabolic activity quenched by freezing samples in dry ice and ethanol, and stored at −80°C. Metabolites were extracted by addition of 600 μl ice-cold 1:1 (vol/vol) methanol/water (containing 1 nmol scyllo-Inositol as internal standard) to the cell pellets, samples were transferred to a chilled microcentrifuge tube containing 300μl chloroform and 600μl methanol (1500 μl total, in 3:1:1 vol/vol methanol/water/chloroform). Samples were sonicated in a water bath for 8 min at 4°C, and centrifuged (13000 rpm) for 10 min at 4°C. The supernatant containing the extract was transferred to a new tube for evaporation in a speed-vacuum centrifuge, resuspended in 3:3:1 (vol/vol/vol) methanol/water/chloroform (350μl total) to phase separate polar metabolites (upper aqueous phase) from apolar metabolites (lower organic phase), and centrifuged. The aqueous phase was transferred to a new tube for evaporation in a speed-vacuum centrifuge, washed with 60 μl methanol, dried again, and derivatized by methoximation (20μl of 20 mg/ml methoxyamine in pyridine, RT overnight) and trimethylsilylation (20μl of N,O-bis(trimetylsilyl)trifluoroacetamide + 1% trimethylchlorosilane) (Sigma, 33148) for ≥ 1 h. GC-MS analysis was performed using an Agilent 7890B-5977A system equipped with a 30 m + 10 m × 0.25 mm DB-5MS + DG column (Agilent J&W) connected to an MS operating in electron-impact ionization (EI) mode. One microliter was injected in splitless mode at 270 °C, with a helium carrier gas. The GC oven temperature was held at 70°C for 2 min and subsequently increased to 295 °C at 12.5 °C/min, then to 320 °C at 25 °C/min (held for 3 min). MassHunter Workstation (B.06.00 SP01, Agilent Technologies) was used for metabolite identification by comparison of retention times, mass spectra and responses of known amounts of authentic standards. Metabolite abundance and mass isotopologue distributions (MID) with correction of natural 13C abundance were determined by integrating the appropriate ion fragments using the GC-MS Assignment Validator and Integrator (GAVIN) (Behrends et al., 2011).

### Quantification of interferon-gamma secretion

Soluble Interferon-gamma (IFN-γ) was quantified in culture media 3 days after CD8+ T cell activation using the Mouse IFN-gamma Quantikine ELISA Kit (RnD Systems, MIF00).

### Oxygen consumption rate and extracellular acidification rate measurements

CD8+ T cells activated for 3 days were washed and oxygen consumption rate (OCR) and extracellular acidification rate (ECAR) in a Seahorse Extracellular Flux Analyzer XF96 (Agilent). 3×10^5^ CD8+ T cells were plated onto poly-D-lysine coated wells and assayed in XF RPMI medium (Agilent) pH 7.4 supplemented with 10 mM glucose and 2 mM glutamine. A minimum of 4 technical replicates per biological replicate was used. During the assay, wells were sequentially injected with 1 μM oligomycin (Sigma), 1.5 μM FCCP (Sigma) and 100 nM rotenone (Sigma) + 1 μM antimycin A (Sigma).

### Glycolytic stress test of CD8+ T cells 12 hours after activation

Purified CD8+ T cells were activated for 12 hours, then 3×10^5^ CD8+ T cells were plated onto poly-D-lysine coated wells in order to form a homogeneous monolayer. Cells were assayed in XF RPMI medium (Agilent) pH 7.4 supplemented with 2 mM glutamine. A minimum of 4 technical replicates per biological replicate was used. Oxygen consumption rate (OCR) and extracellular acidification rate (ECAR) were determined through a Seahorse Extracellular Flux Analyzer XF96 (Agilent). During the assay, wells were sequentially injected with 40 mM NaCl/NaLac with or without the MCT1-selective inhibitor AZD3965 (Cayman Chemical, 19912) or the LDH inhibitor GSK2837808A (Bio-Techne, 5189), or plain media alone, followed by 10 mM glucose (Sigma), 1 μM oligomycin and 50 mM 2-DG.

### Statistics

Statistical analyses were performed with Prism 9 version 9.2.0 (GraphPad). Statistical tests and number of replicates are stated in figure legends.

### Study approval

All animal experiments were approved by the regional animal ethics Committee of Northern Stockholm, Sweden (dnr N78/15 and N101/16). The Stockholm Regional Ethical Review Board does not require an ethical permission for the use of nonidentified healthy donor blood samples.

## Acknowledgements

The authors acknowledge Dr. Jernej Ule (Francis Crick Institute) for advice and resources on metabolomic experiments; Dr. David Macias (University of Cambridge) for advice on osmolality measurements; Dr. Jeffrey E. Mold (Karolinska Institutet) for advice on human T cell experiments. Metabolomics experiments were performed at the Swedish Metabolomics Center (Umeå, Sweden). [U-^13^C_6_]glucose metabolomics experiments were performed in the Metabolomics STP at The Francis Crick Institute (London, UK), which receives its core funding from Cancer Research UK (FC001999), the UK Medical Research Council (FC001999), and the Wellcome Trust (FC001999). RNA-sequencing and bioinformatic analysis was performed at the Bioinformatics and Expression Analysis (BEA) Core Facility at the Department of Biosciences and Nutrition, which is supported by the board of research at the Karolinska Institute and the research committee at the Karolinska hospital. The work was funded by the Knut and Alice Wallenberg Scholar Award, the Swedish Medical Research Council (Vetenskapsrådet 2019-01485), the Swedish Cancer Fund (Cancerfonden, CAN2018/808), the Swedish Children’s Cancer Fund (Barncancerfonden PR2020-007), the Portuguese Foundation for Science and Technology scholarship to P.P. Cunha (SFRH/BD/115612/2016), and the Principal Research Fellowship (214283/Z/18/Z) to Randall S. Johnson from the Wellcome Trust.

## Competing interests

The authors declare that no competing interests exist

## Additional files

### Supplementary files

Transparent reporting form

### Data availability

All data generated are included in the manuscript and supporting files.

The RNA sequencing data discussed in this publication have been deposited in NCBI’s Gene

Expression Omnibus (Edgar et al., 2002) and are accessible through GEO Series accession number GSE190808 (https://www.ncbi.nlm.nih.gov/geo/query/acc.cgi?acc=GSE190808).”

## Supplementary Figures

**Supplementary Figure 1.**
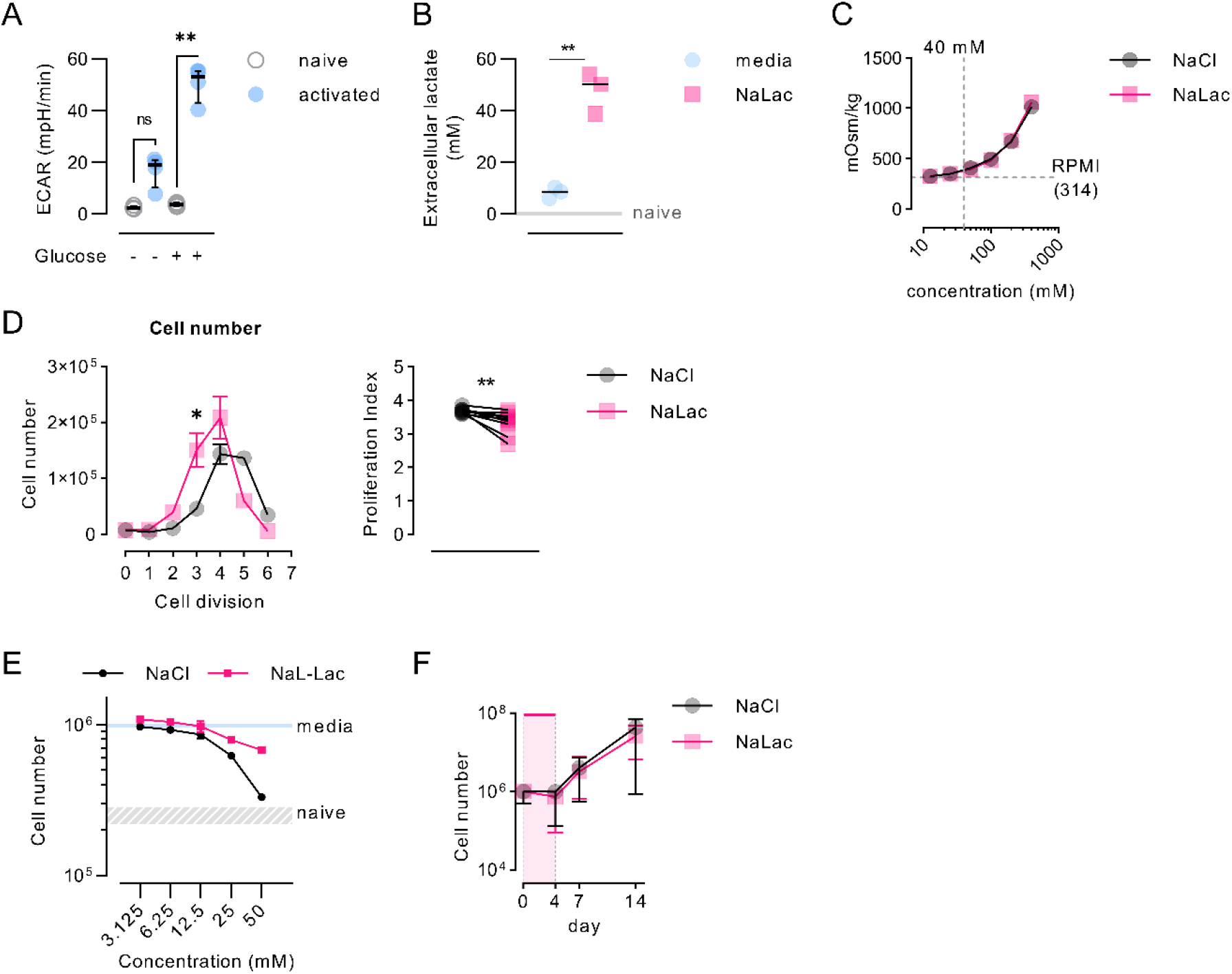
Sodium lactate, but not lactic acid, is well-tolerated by CD8+ T cells. **(A)** Mouse CD8+ T cells were activated for 72h with anti-CD3/CD28 beads or kept in a naïve-like state (non-activated) with IL-7 over the same period of time. After activation, cells were analyzed in a Seahorse XF96 Flux Analyzer to test their glycolytic function, via consecutive injection of 10 mM glucose (glc), 1 μM oligomycin (oligo) and 50 mM 2-deoxyglucose (2-DG). ECAR: Extracellular Acidification Rate. Data are the median and interquartile range of n = 4 independent mouse donors. ** p<0.005 oneway ANOVA with Tukey’s multiple comparisons test. **(B)** Extracellular lactate concentration from the supernatant of mouse CD8+ T cells, 72 hours after *in vitro* activation. Lactate levels were measured by enzymatic assay. Dotted line: non-activated (naïve) cells. N = 3 individual mice. ** P<0.01 paired t test, two-tailed. **(C)** Osmolality of sodium chloride (NaCl) and sodium lactate (NaLac) solutions in complete RPMI. Horizontal dotted line: RPMI reference. Vertical dotted line: osmolality of 40 mM NaCl/NaLac solution in RPMI. **(D)** Mouse CD8+ T cells were activated for 72 hours in the presence of 40 mM NaCl or NaLac. At the end of the activation period, cells were washed and absolute cell number and proliferation index at each cell generation were determined by CFSE dilution. Mean and SEM of n = 3 independent mouse donors. * P<0.001 repeated-measures two-way ANOVA with Sidak’s multiple comparison test. **(E)** Effect of varying concentrations of NaCl or NaLac on human CD8+ T cell number, 96 hours after activation with anti-CD3/CD28 beads and in the presence of IL-2. Dotted line and grey range: average of non-activated (naive) cells. Solid light blue line: cells activated in the presence of media (control). N = 2 independent human donors. **(F)** Human CD8+ T cells purified from healthy donor PMBCs were activated in the presence of 40 mM NaCl or NaLac for 4 days. Long-term expansion was continued after NaCl/NaLac removal at day 4, with addition of fresh media and IL-2. Cell number was measured at day 4, 7 and 14. Data are the median and interquartile range of n = 10 human donors.

**Supplementary Figure 2.**
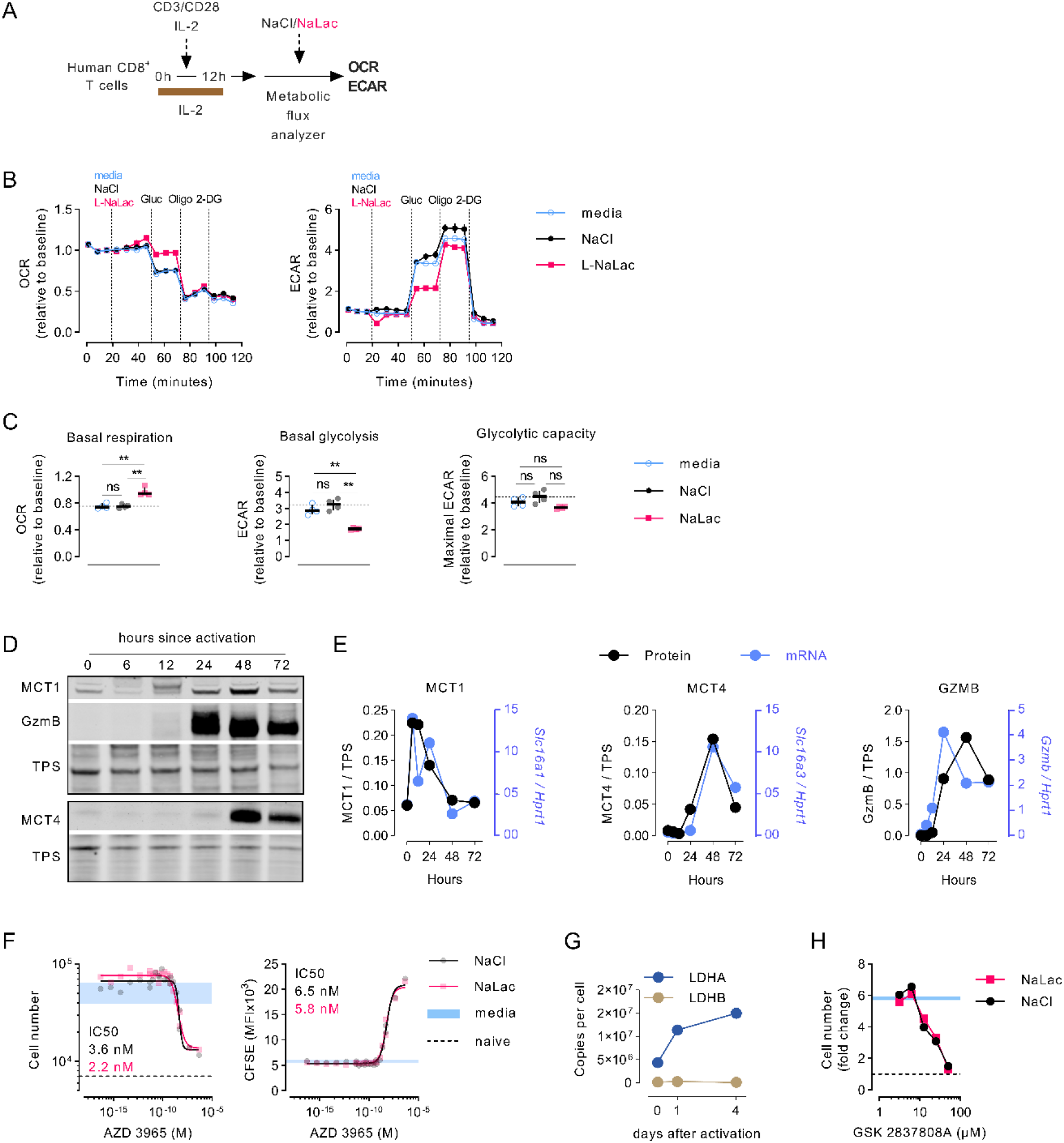
Lactate modulates CD8+ T cell metabolism. **(A)** Experimental design. Human CD8+ T cells were isolated from the blood of healthy donors and activated for 12 hours in IL-2 and. After 12 hours, cells were transferred in a seahorse XF analyzer for metabolic flux analysis after injection of 40 mM NaCl or sodium lactate (NaLac). **(B)** Glycolytic stress test of human CD8+ T cells 12 hours after activation. Graphs show extracellular acidification rate (ECAR) and oxygen consumption rate (OCR) during sequential injection of either media, or 40 mM NaCl, or 40 mM NaLac, followed by 10 mM glucose, 1 μM oligomycin (oligo) and 50 mM 2-deoxyglucose (2-DG). ECAR and OCR are normalized to baseline levels. Data is the mean and SEM of 4 technical replicates of one representative donor. **(C)** Basal glycolysis, basal mitochondrial respiration, and glycolytic capacity determined from the assay shown in (B). Lines indicate the median of n = 4 technical replicates of one representative donor. Dotted line: median of NaCl-treated cells. ** P<0.01 one-way ANOVA with Holm-Šídák’s multiple comparisons test between each donor’s technical replicates. **(D)** 15 μg of mouse CD8+ T cell protein cytoplasmic extracts from 0, 6, 12, 24, 48, and 72 hours post-activation were probed with antibodies against MCT1, MCT4, and GZMB. Total protein stain (TPS) was used as loading control. Data is provided as Figure 2-figure supplement-source data 1. **(E)** Protein quantification from experiment shown in (D) (normalized to total protein stain, TPS) and mRNA (normalized to expression of the housekeeping gene *Hprt*) levels of MCT1, MCT4 and GZMB over time since activation. **(F)** Mouse CD8+ T cells were activated for 72 hours in the presence of the monocarboxylate transporter-1 (MCT1) inhibitor AZD3985 in combination with 40 mM NaLac or plain media. A non-linear fit ([Inhibitor] vs. response -- Variable slope (four parameters)) was used to determine IC_50_. Blue area = range of untreated controls; dotted line: non-activated (naive) cells. **(G)** Estimated protein mean copy number of Ldha and Ldhb at day 0, 1, and 4 after activation of mouse P14 transgenic CD8+ T cells in the presence of IL-2. Data were obtained from the public ImmPRes database (http://immpres.co.uk/) (Howden et al., 2019). Lines represent the mean and standard deviation. **(H)** CD8+ T cells were activated for 72 hours in the presence of increasing doses of the LDH inhibitor GSK2837808A and either 40 mM NaCl or NaLac. Cell numbers are expressed as fold change relative to seeding cell number.

**Supplementary Figure 3.**
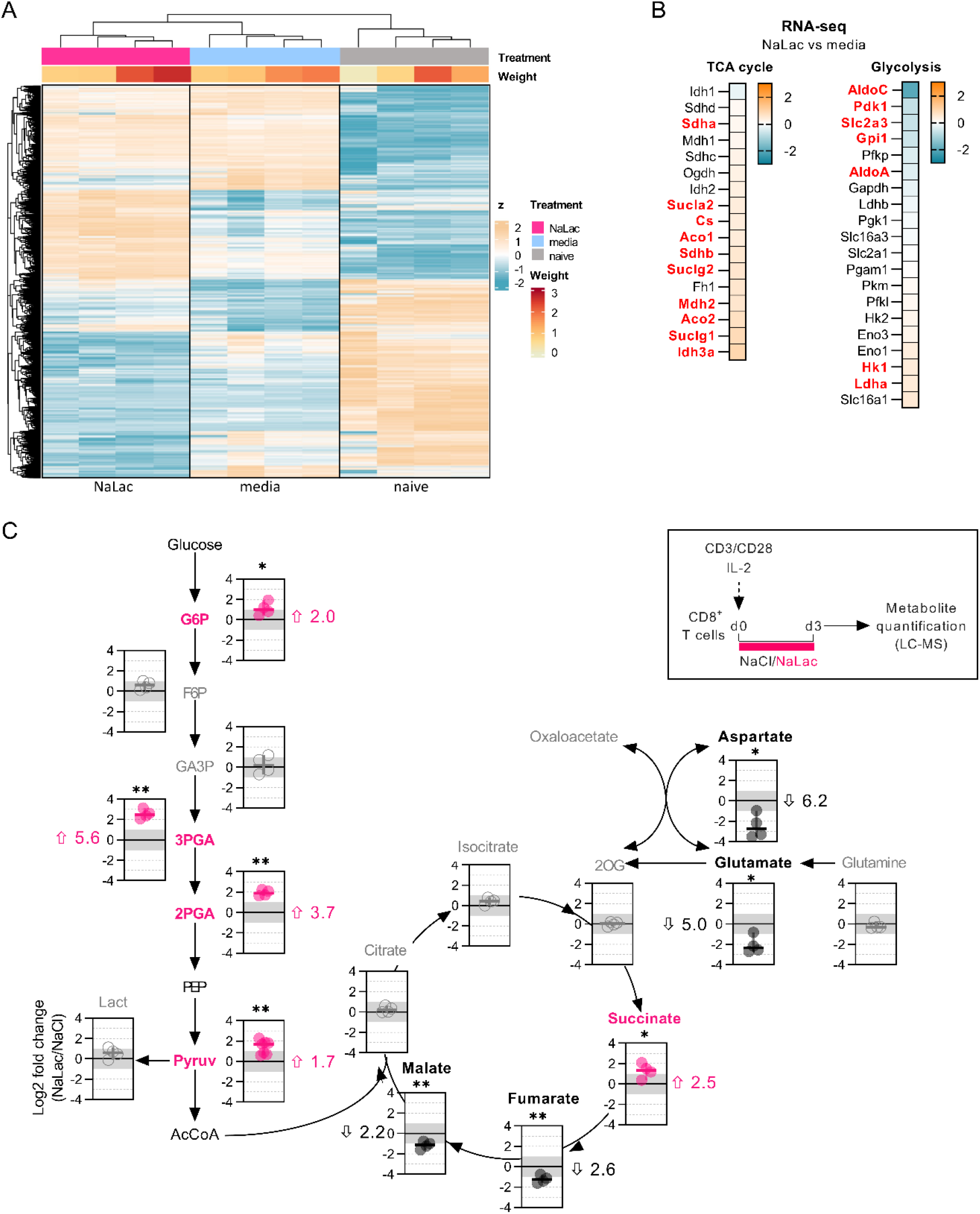
Early exposure to lactate alters CD8+ T cell metabolism and gene expression. **(A)** Heatmap and clustering of genes from RNA-seq experiment described in Figure 3A. Only genes that were found significantly deregulated in any of the comparisons are shown. Each gene (row) is standardized (z) to a mean of 0 and standard deviation (SD) of 1. “Treatment” indicates activation in the presence of either media (light blue) or NaLac (pink), or non-activated cells (naive, grey). The “weight” heatmap indicates the degree of influence of each sample on the overall analysis. Data is provided as Figure 3-figure supplement-source data 1. **(B)** Same data as in Figure 3C, shown as heatmaps of the differentially expressed TCA cycle (left) or glycolytic (right) genes, in NaLac-relative to media-treated CD8+ T cells. Colors indicate the degree of differential expression (as Log_2_ of the fold change) of genes in each pathway. Significant genes (adjPval ≤ 0.01) are indicated in red. Data is provided as Figure 3-figure supplement-source data 2. **(C)** Experimental design and metabolic map of the same experiment shown in Figure 3D. Mouse CD8+ T cells were activated for 72 hours in the presence of 40 mM NaCl or NaLac. Metabolic changes after activation were quantified by sugar phosphate and GC-MS analysis. Data show log_2_ fold changes between NaLac- and NaCl-treated samples of n = 4 independent mouse donors. Pink and black data points represent significantly increased or decreased metabolite frequency in NaLac versus NaCl-treated cells, respectively. Median fold-change is indicated next to each plot. ** P<0.01, *P<0.05, two-tailed ratio paired t-test. Data is provided as Figure 3-figure supplement-source data 3.

**Supplementary Figure 4.**
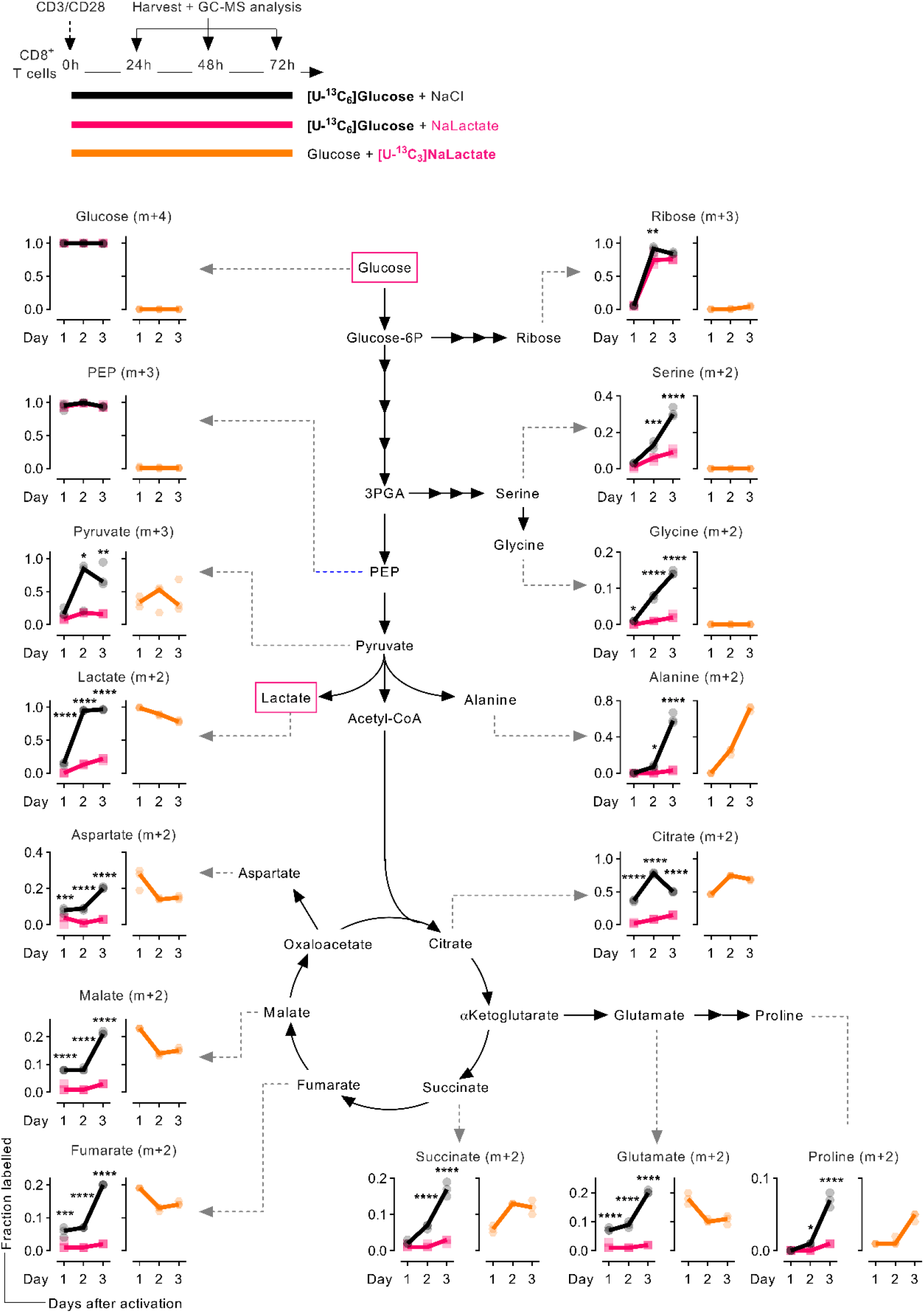
Lactate is incorporated in cellular metabolism and displaces glucose. Same experiment as in Figure 4, but shown as metabolic diagrams of glucose- or lactate-derived carbons incorporation into cellular metabolites at 24, 48, and 72 hours after activation. Top: Experimental design and schematic representation of the labelled molecules. Bottom: lines showing the fraction of each metabolite labelled by [U-^3^C_6_]glucose alone (black), or in the presence of sodium lactate (pink), or by [U-^3^C_6_]sodium lactate (orange). Lines connect the median of n = 3 independent mice donors. * P<0.05, ** P<0.01, *** P<0.001, repeated-measures two-way ANOVA with Sidak’s multiple comparison test comparing glucose alone and glucose with addition of 40 mM NaLac.

**Supplementary Figure 5.**
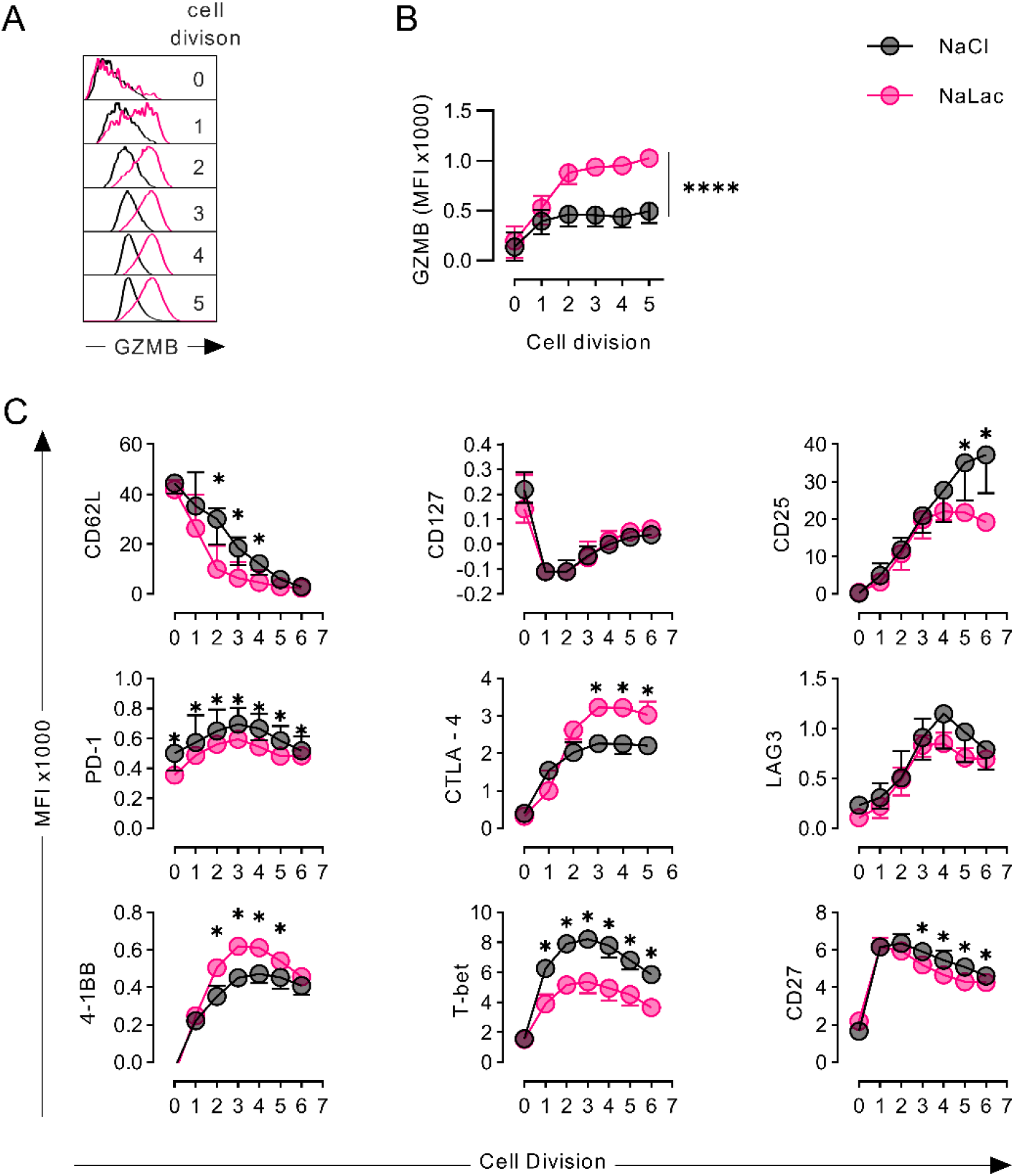
Activation in the presence of lactate modulates the differentiation of CD8+ T cells. **(A)** Mouse CD8+ T cells were activated for 72 hours in the presence of either 40 mM NaCl (black line) or 40 mM NaLac (pink line). Flow cytometry histograms showing intracellular Granzyme B (GZMB) levels at each cell division (determined by CTV peaks). **(B)** Median Fluorescence Intensity (MFI) of GZMB levels at each cell division of mouse CD8+ T cells activated as in (A). Lines represent the median of n = 6 biological replicates. **** P<0.0001 Two-way Anova with Šídák’s multiple comparisons test. **(C)** Median Fluorescence Intensity of CD62L, CD127, CD25, PD-1, CTLA-4, LAG-3, 4-1BB, T-bet, and CD27 at each cell division, 72 hours after activation with 40 mM NaCl (black line) or NaLac (pink line). Significance was calculated using RM two-way ANOVA with the Geisser-Greenhouse correction and Sidak’s multiple comparison test.

## Notes

### Competing Interest Statement

The authors have declared no competing interest.

## References

Ashley Menk A v, Scharping NE, Moreci RS, Young HA, Gelhaus Wendell S, Delgoffe Correspondence GM. 2018. Early TCR Signaling Induces Rapid Aerobic Glycolysis Enabling Distinct Acute T Cell Effector Functions. Cell Reports 22. doi:10.1016/j.celrep.2018.01.040

Baltazar F, Afonso J, Costa M, Granja S. 2020. Lactate Beyond a Waste Metabolite: Metabolic Affairs and Signaling in Malignancy. Frontiers in Oncology 0:231. doi:10.3389/FONC.2020.00231

Behrends V, Tredwell GD, Bundy JG. 2011. A software complement to AMDIS for processing GC-MS metabolomic data. Analytical Biochemistry 415:206–208. doi:10.1016/J.AB.2011.04.009

Billiard J, Dennison JB, Briand J, Annan RS, Chai D, Colón M, Dodson CS, Gilbert SA, Greshock J, Jing J, Lu H, McSurdy-Freed JE, Orband-Miller LA, Mills GB, Quinn CJ, Schneck JL, Scott GF, Shaw AN, Waitt GM, Wooster RF, Duffy KJ. 2013. Quinoline 3-sulfonamides inhibit lactate dehydrogenase A and reverse aerobic glycolysis in cancer cells. Cancer & metabolism 1. doi:10.1186/2049-3002-1-19

Brand A, Singer K, Koehl GE, Kolitzus M, Schoenhammer G, Thiel A, Matos C, Bruss C, Klobuch S, Peter K, Kastenberger M, Bogdan C, Schleicher U, Mackensen A, Ullrich E, Fichtner-Feigl S, Kesselring R, Mack M, Ritter U, Schmid M, Blank C, Dettmer K, Oefner PJ, Hoffmann P, Walenta S, Geissler EK, Pouyssegur J, Villunger A, Steven A, Seliger B, Schreml S, Haferkamp S, Kohl E, Karrer S, Berneburg M, Herr W, Mueller-Klieser W, Renner K, Kreutz M. 2016. LDHA-Associated Lactic Acid Production Blunts Tumor Immunosurveillance by T and NK Cells. Cell Metabolism 24:657–671. doi:10.1016/j.cmet.2016.08.011

Brooks GA. 2020. Lactate as a fulcrum of metabolism. Redox Biology. doi:10.1016/j.redox.2020.101454

Brooks GA. 2018. The Science and Translation of Lactate Shuttle Theory. Cell Metabolism. doi:10.1016/j.cmet.2018.03.008

Carr EL, Kelman A, Wu GS, Gopaul R, Senkevitch E, Aghvanyan A, Turay AM, Frauwirth KA. 2010. Glutamine Uptake and Metabolism Are Coordinately Regulated by ERK/MAPK during T Lymphocyte Activation. The Journal of Immunology 185:1037–1044. doi:10.4049/jimmunol.0903586

Chen YJ, Mahieu NG, Huang X, Singh M, Crawford PA, Johnson SL, Gross RW, Schaefer J, Patti GJ. 2016. Lactate metabolism is associated with mammalian mitochondria. Nature Chemical Biology 12:937–943. doi:10.1038/nchembio.2172

Chih-Hao Chang, Erika L Pearce. 2016. Emerging concepts of T cell metabolism as a target of immunotherapy. Nature immunology 17:364–368. doi:10.1038/NI.3415

Chih-Hao Chang, Jing Qiu, David O’Sullivan, Michael D. Buck, Takuro Noguchi, Jonathan D. Curtis, Qiongyu Chen, Mariel Gindin, Matthew M. Gubin, Gerritje J.W. van der Windt, Elena Tonc, Robert D. Schreiber, Edward J. Pearce, Erika L. Pearce. 2015. Metabolic Competition in the Tumor Microenvironment Is a Driver of Cancer Progression. Cell 162:1229–1241. doi:10.1016/J.CELL.2015.08.016

Dobin A, Davis CA, Schlesinger F, Drenkow J, Zaleski C, Jha S, Batut P, Chaisson M, Gingeras TR. 2013. STAR: ultrafast universal RNA-seq aligner. Bioinformatics 29:15. doi:10.1093/BIOINFORMATICS/BTS635

Doedens AL, Phan AT, Stradner MH, Fujimoto JK, Nguyen J v., Yang E, Johnson RS, Goldrath AW. 2013. Hypoxia-inducible factors enhance the effector responses of CD8 + T cells to persistent antigen. Nature Immunology 14:1173–1182. doi:10.1038/ni.2714

Faubert B, Li KY, Cai L, Hensley CT, Kim J, Zacharias LG, Yang C, Do QN, Doucette S, Burguete D, Li H, Huet G, Yuan Q, Wigal T, Butt Y, Ni M, Torrealba J, Oliver D, Lenkinski RE, Malloy CR, Wachsmann JW, Young JD, Kernstine K, DeBerardinis RJ. 2017. Lactate Metabolism in Human Lung Tumors. Cell 171:358–371.e9. doi:10.1016/j.cell.2017.09.019

Finlay DK, Rosenzweig E, Sinclair L v., Carmen FC, Hukelmann JL, Rolf J, Panteleyev AA, Okkenhaug K, Cantrell DA. 2012. PDK1 regulation of mTOR and hypoxia-inducible factor 1 integrate metabolism and migration of CD8+ T cells. Journal of Experimental Medicine 209:2441–2453. doi:10.1084/jem.20112607

Fischer K, Hoffmann P, Voelkl S, Meidenbauer N, Ammer J, Edinger M, Gottfried E, Schwarz S, Rothe G, Hoves S, Renner K, Timischl B, Mackensen A, Kunz-Schughart L, Andreesen R, Krause SW, Kreutz M. 2007. Inhibitory effect of tumor cell-derived lactic acid on human T cells. Blood 109:3812–3819. doi:10.1182/blood-2006-07

Grist JT, Jarvis LB, Georgieva Z, Thompson S, Sandhu HK, Burling K, Clarke A, Jackson S, Wills M, Gallagher FA, Jones JL. 2018. Extracellular Lactate: A Novel Measure of T Cell Proliferation. The Journal of Immunology Author Choice 200:1220. doi:10.4049/JIMMUNOL.1700886

Gullberg J, Jonsson P, Nordström A, Sjöström M, Moritz T. 2004. Design of experiments: an efficient strategy to identify factors influencing extraction and derivatization of Arabidopsis thaliana samples in metabolomic studies with gas chromatography/mass spectrometry. Analytical biochemistry 331:283–295. doi:10.1016/J.AB.2004.04.037

Haas R, Smith J, Rocher-Ros V, Nadkarni S, Montero-Melendez T, D’Acquisto F, Bland EJ, Bombardieri M, Pitzalis C, Perretti M, Marelli-Berg FM, Mauro C. 2015. Lactate regulates metabolic and proinflammatory circuits in control of T cell migration and effector functions. PLoS Biology 13:1–24. doi:10.1371/journal.pbio.1002202

Halestrap AP, Meredith D. 2003. The SLC16 gene family - From monocarboxylate transporters (MCTs) to aromatic amino acid transporters and beyond. European Journal of Physiology. doi:10.1007/s00424-003-1067-2

Halestrap AP, Price NT. 1999. The proton-linked monocarboxylate transporter (MCT) family: Structure, function and regulation. Biochemical Journal. doi:10.1042/0264-6021:3430281

Hayashi K, Jutabha P, Endou H, Sagara H, Anzai N. 2013. LAT1 Is a Critical Transporter of Essential Amino Acids for Immune Reactions in Activated Human T Cells. The Journal of Immunology 191:4080–4085. doi:10.4049/jimmunol.1300923

Heiden MGV, Cantley LC, Thompson CB. 2009. Understanding the warburg effect: The metabolic requirements of cell proliferation. Science 324:1029–1033. doi:10.1126/science.1160809

Howden AJM, Hukelmann JL, Brenes A, Spinelli L, Sinclair L v., Lamond AI, Cantrell DA. 2019. Quantitative analysis of T cell proteomes and environmental sensors during T cell differentiation. Nature Immunology 2019 20:11 20:1542–1554. doi:10.1038/s41590-019-0495-x

Hui S, Ghergurovich JM, Morscher RJ, Jang C, Lu W, Esparza LA, Reya T, Zhan L, Guo Y, White E, Rabinowitz JD. 2018. Glucose feeds the TCA cycle via circulating lactate. Nature 551:115–118. doi:10.1038/nature24057.Glucose

Hukelmann JL, Anderson KE, Sinclair L v, Grzes KM, Murillo AB, Hawkins PT, Stephens LR, Lamond AI, Cantrell DA. 2016. The cytotoxic T cell proteome and its shaping by the kinase mTOR. Nature immunology 17:104–12. doi:10.1038/ni.3314

Jonathan W. Heusel, Robin L. Wesselschmidt, Sujan Shresta, John H. R, Timothy J. Ley. 1994. Cytotoxic lymphocytes require granzyme B for the rapid induction of DNA fragmentation and apoptosis in allogeneic target cells. Cell 76:977–987. doi:10.1016/0092-8674(94)90376-X

Klein Geltink RI, Kyle RL, Pearce EL. 2018. Unraveling the Complex Interplay Between T Cell Metabolism and Function. Annual Rev Immunol 36:461–488. doi:10.1146/annurev-immunol-042617-053019

Liu R, Holik AZ, Su S, Jansz N, Chen K, Leong HS, Blewitt ME, Asselin-Labat ML, Smyth GK, Ritchie ME. 2015. Why weight? Modelling sample and observational level variability improves power in RNA-seq analyses. Nucleic acids research 43. doi:10.1093/NAR/GKV412

Ma EH, Bantug G, Griss T, Richer MJ, Hess C, Jones Correspondence RG. 2017. Serine Is an Essential Metabolite for Effector T Cell Expansion. Cell Metabolism 25:345–357. doi:10.1016/j.cmet.2016.12.011

Ma EH, Verway MJ, Johnson RM, Roy DG, Steadman M, Hayes S, Williams KS, Sheldon RD, Samborska B, Kosinski PA, Kim H, Griss T, Faubert B, Condotta SA, Krawczyk CM, DeBerardinis RJ, Stewart KM, Richer MJ, Chubukov V, Roddy TP, Jones RG. 2019. Metabolic Profiling Using Stable Isotope Tracing Reveals Distinct Patterns of Glucose Utilization by Physiologically Activated CD8+ T Cells. Immunity 51:856–870.e5. doi:10.1016/j.immuni.2019.09.003

Manley G, Conn JR, Catchpoole EM, Runnegar N, Mapp SJ, Markey KA. 2013. The mTORC1 pathway stimulates glutamine metabolism and cell proliferation by repressing SIRT4. Cell 32:736–740. doi:10.1016/j.cell.2013.04.023.The

Mendler AN, Hu B, Prinz PU, Kreutz M, Gottfried E, Noessner E. 2012. Tumor lactic acidosis suppresses CTL function by inhibition of p38 and JNK/c-Jun activation. International Journal of Cancer 131:633–640. doi:10.1002/ijc.26410

O’Neill LAJ, Kishton RJ, Rathmell J. 2016. A guide to immunometabolism for immunologists. Nature Reviews Immunology 16:553–565. doi:10.1038/nri.2016.70

Palazon A, Goldrath AW, Nizet V, Johnson RS. 2014. HIF Transcription Factors, Inflammation, and Immunity. Immunity 41:518–528. doi:10.1016/j.immuni.2014.09.008

Palazon A, Tyrakis PA, Macias D, Goldrath AW, Bergh J, Correspondence RSJ. 2017. An HIF-1a/VEGF-A Axis in Cytotoxic T Cells Regulates Tumor Progression Highlights d HIF-1a drives CD8 + T cell migration and effector function. doi:10.1016/j.ccell.2017.10.003

Pearce EL. 2010. Metabolism in T cell activation and differentiation. current opinion in immunology 22:314–320. doi:10.1016/j.coi.2010.01.018

Pearce EL, Poffenberger MC, Chang C-H, Jones RG. 2013. Fueling Immunity: Insights into Metabolism and Lymphocyte Function. Science 342:1242454–1242454. doi:10.1126/science.1242454

Péronnet F, Aguilaniu B. 2006. Lactic acid buffering, nonmetabolic CO2 and exercise hyperventilation: A critical reappraisal. Respiratory Physiology and Neurobiology. doi:10.1016/j.resp.2005.04.005

Ping-Chih Ho, Jessica Dauz Bihuniak, Andrew N. Macintyre, Matthew Staron, Xiaojing Liu, Robert Amezquita, Yao-Chen Tsui, Guoliang Cui, Goran Micevic, Jose C. Perales, Steven H. Kleinstein, E. Dale Abel, Karl L. Insogna, Stefan Feske, Jason W. Locasale, Marcus W. Bosenberg, Jeffrey C. Rathmell, Susan M. Kaech. 2015. Phosphoenolpyruvate Is a Metabolic Checkpoint of Antitumor T Cell Responses. Cell 162:1217–1228. doi:10.1016/J.CELL.2015.08.012

Robert A Robergs, Farzenah Ghiasvand, Daryl Parker. 2004. Biochemistry of exercise-induced metabolic acidosis. American journal of physiology Regulatory, integrative and comparative physiology 287. doi:10.1152/AJPREGU.00114.2004

Robinson MD, McCarthy DJ, Smyth GK. 2010. edgeR: a Bioconductor package for differential expression analysis of digital gene expression data. Bioinformatics (Oxford, England) 26:139–140. doi:10.1093/BIOINFORMATICS/BTP616

Rundqvist H, Velica P, Barbieri L, Gameiro PA, Bargiela D, Gojkovic M, Mijwel S, Reitzner SM, Johnson RS. 2020. Cytotoxic T-cells mediate exercise-induced reductions in tumor growth. eLife 2020;9:e59996 1–25. doi:DOI: https://doi.org/10.7554/eLife.59996

Tracy S P Heng, Michio W Painter, The Immunological Genome Project Consortium. 2008. The Immunological Genome Project: networks of gene expression in immune cells. Nature Immunology 9:1091–1094.

Tyrakis PA, Palazon A, Macias D, Lee KL, Phan AT, Veliça P, You J, Chia GS, Sim J, Doedens A, Abelanet A, Evans CE, Griffiths JR, Poellinger L, Goldrath AW, Johnson RS. 2016. S-2-hydroxyglutarate regulates CD8+ T-lymphocyte fate. Nature 540:236–241. doi:10.1038/nature20165

Wang R, Dillon CP, Shi LZ, Milasta S, Carter R, Finkelstein D, McCormick LL, Fitzgerald P, Chi H, Munger J, Green DR. 2011. The Transcription Factor Myc Controls Metabolic Reprogramming upon T Lymphocyte Activation. Immunity 35:871–882. doi:10.1016/j.immuni.2011.09.021

Wang R, Green DR. 2012. Metabolic reprogramming and metabolic dependency in T cells. Immunological Reviews 249:14–26. doi:10.1111/j.1600-065X.2012.01155.x

Watson MLJ, Vignali PDA, Mullett SJ, Overacre-Delgoffe AE, Peralta RM, Grebinoski S, Menk A v., Rittenhouse NL, DePeaux K, Whetstone RD, Vignali DAA, Hand TW, Poholek AC, Morrison BM, Rothstein JD, Wendell SG, Delgoffe GM. 2021. Metabolic support of tumour-infiltrating regulatory T cells by lactic acid. Nature 2021 591:7851 591:645–651. doi:10.1038/s41586-020-03045-2

Zhang D, Tang Z, Huang H, Zhou G, Cui C, Weng Y, Liu W, Kim S, Lee S, Perez-Neut M, Ding J, Czyz D, Hu R, Ye Z, He M, Zheng YG, Shuman HA, Dai L, Ren B, Roeder RG, Becker L, Zhao Y. 2019. Metabolic regulation of gene expression by histone lactylation. Nature 574:575–580. doi:10.1038/s41586-019-1678-1

